# Dual-Antigen CCHFV Vaccine Induces Neutralizing Immunity and Protection Against Lethal Challenge

**DOI:** 10.64898/2026.10.04.756518

**Authors:** Jingyi Yan, Lara Sweetapple, Christian An Bính Nordentoft, Sofia Appelberg, Stéphanie Devignot, Vanessa Monteil, Pia Binder, Michaela Mochánová, Elias Obinger, Drew Weissman, Norbert Pardi, Lars Frelin, Ali Mirazimi, Matti Sällberg, Gustaf Ahlén

**Affiliations:** Department of Laboratory Medicine, Karolinska Institutet, Stockholm, Sweden; Clinical Microbiology, Karolinska University Hospital, Stockholm, Sweden; Karolinska Advanced Therapy Medicinal Products (ATMP) Center, Stockholm, Sweden; Public Health Agency of Sweden, Stockholm, Sweden; Department of Pharmacy, Faculty of Pharmacy, Uppsala University, Uppsala, Sweden; Department of Gene Technology, KTH Royal Institute of Technology, Stockholm, Sweden; Department of Microbiology, Perelman School of Medicine, University of Pennsylvania, Philadelphia, PA, USA; Medical Diagnostics Karolinska, Karolinska University Hospital, Stockholm, Sweden

**Keywords:** Crimean-Congo hemorrhagic fever virus (CCHFV), DNA vaccine, mRNA vaccine, in vivo Electroporation, Neutralizing antibodies, Immunogenicity, Lipid nanoparticle (LNP), Nucleoprotein (NP), Glycoprotein Gc

## Abstract

Crimean-Congo hemorrhagic fever virus (CCHFV) is a WHO-priority pathogen with high mortality and wide global distribution. A major challenge in CCHFV vaccine development is identifying antigens that induce functional cellular and humoral immunity. We previously showed that the CCHF nucleoprotein (NP) elicits protective responses in mice and non-human primates but does not induce neutralizing antibodies. We therefore systematically screened six chimeric DNA vaccines expressing different CCHF glycoproteins combined with NP. A construct comprising the ectodomain of the glycoprotein Gc and NP elicited robust antibody and T-cell responses to both antigens. The Gc-NP gene was subsequently developed as a lipid nanoparticle-formulated mRNA vaccine using a scalable manufacturing pipeline. Gc-NP mRNA-LNP induced robust antigen-specific humoral and cellular responses comparable to those elicited by the corresponding DNA vaccine, despite the substantially lower dose. Both vaccine approaches generated Gc-directed neutralizing antibody responses against CCHFV pseudovirus and protected mice against homologous Hoti and heterologous IbAr10200 strain challenge. These findings demonstrate that systematic antigen screening combined with genetic vaccine platforms can identify a potent multi-antigen CCHF vaccine candidate.

**HIGHLIGHTS:**

1. A systematic screening of DNA-based genes identified a highly immunogenic combination of the Gc protein ectodomain and NP.
2. This novel vaccine candidate induced Gc-specific neutralizing antibodies and T-cell responses against both NP and Gc.
3. An mRNA-LNP vaccine expressing the Gc-NP gene further improved immunogenicity and conferred protection against lethal challenge with both CCHFV Hoti and IbAr10200 strains in mice.

**GRAPHICAL ABSTRACT:** 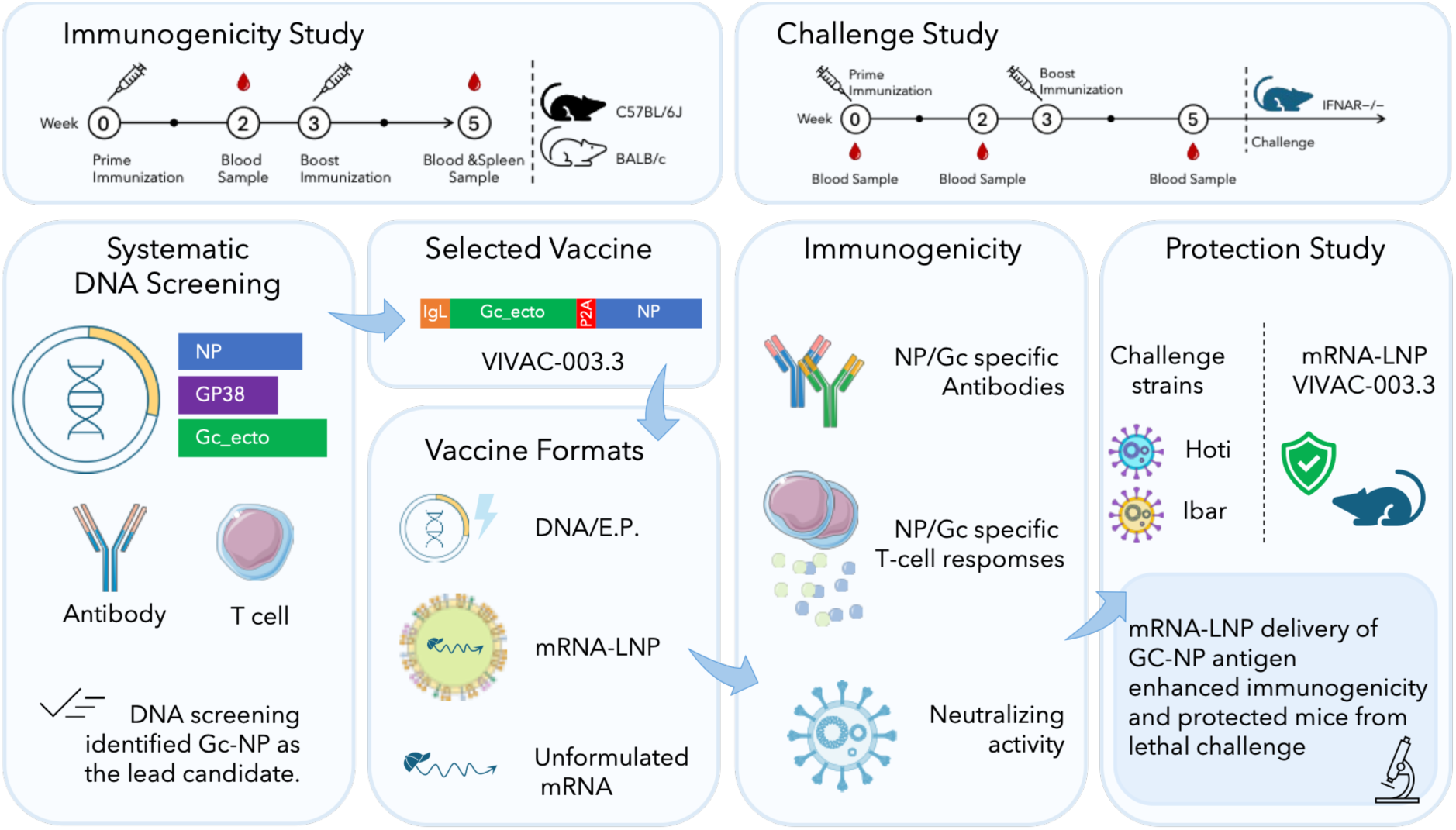

## INTRODUCTION

Crimean-Congo hemorrhagic fever (CCHF) is a severe tick-borne disease in humans with case fatality rates of up to ∼30%. The causative agent, CCHF virus (CCHFV), is widely distributed across Africa, Southern and Eastern Europe, the Middle East, and Asia, reflecting the geographic range of Hyalomma tick vectors ^1,2^. Human infection occurs primarily through tick bites, contact with blood or tissues of infected livestock, or exposure to blood and bodily fluids from infected patients, placing rural populations, livestock workers, and health care personnel at increased risk ^1,3,4^. Clinically, CCHF typically presents as an acute, nonspecific febrile illness with thrombocytopenia, hepatic injury, and coagulation abnormalities ^5^, but severe disease may rapidly progress to systemic inflammation, hemorrhage, disseminated intravascular coagulation, multiorgan failure, shock, and death ^6,7^. There is currently no globally approved vaccine against CCHF, and clinical management remains largely supportive. Ribavirin may provide benefit when administered early, whereas evidence for favipiravir remains predominantly preclinical ^8,9^. Early diagnosis is often challenging because the initial clinical manifestations are nonspecific and may overlap with those of other acute febrile illnesses, thereby narrowing the window for timely intervention ^9^. In the absence of broadly approved vaccines and effective countermeasures, CCHFV represents a substantial and persistent public health threat with significant epidemic potential ^10^. Accordingly, the World Health Organization has classified CCHFV as a priority disease for research and development efforts, underscoring the need for vaccine strategies capable of inducing rapid, robust, and durable immunity ^10–12^.

CCHFV is an enveloped, negative-sense RNA virus of the family *Nairoviridae* with a tripartite genome encoding the RNA-dependent RNA polymerase (L segment), nucleoprotein (NP; S segment), and glycoprotein precursor (M segment) processed into the Gn, Gc, and GP38 proteins ^1^. Although CCHFV exhibits substantial genetic diversity ^13^, conserved or functionally constrained epitopes within NP ^14,15^ and Gc ^17–19^ provide a rationale for vaccine strategies aimed at eliciting immune responses with potential cross-strain reactivity ^14,15,19^. The Kosova Hoti (Hoti) strain was selected as the antigenic basis for vaccine design because its complete genome has been well characterized ^20^; Hoti-based infection models in immunocompetent mice and nonhuman primates recapitulate key features of human CCHF ^21,22^. Hoti-derived NP and glycoprotein antigens have demonstrated cross-strain immunogenicity and protection in heterologous CCHFV challenge models ^14–16,23^.

In this study, we developed a panel of CCHFV vaccine candidates using a mechanistically guided antigen-selection strategy aimed at balancing humoral and cellular immunity. Conserved Hoti-derived antigens were selected to promote cross-reactive immune responses, with constructs incorporating the nucleoprotein (NP) and the Gc ectodomain, the latter used in place of the full glycoprotein precursor to focus immune responses on neutralization-relevant epitopes and reduce reliance on complex intracellular processing. The non-structural antigen GP38 with variable immune protection was included in the screening ^24,25^. Candidate constructs were expressed individually or in defined combinations and initially screened using DNA vaccination delivered by *in vivo* electroporation, enabling efficient prioritization based on antibody responses and T-cell responses. The lead NP-Gc construct identified through this multiparametric screening approach was subsequently reformulated as a lipid nanoparticle (LNP)-encapsulated mRNA vaccine and evaluated in comparative immunogenicity studies alongside electroporated DNA and unformulated mRNA, followed by protection against lethal challenge with both the homologous Hoti strain and the heterologous IbAr strain of CCHFV in mice. In parallel, an internal scalable mRNA-LNP manufacturing workflow was established to support rapid candidate translational advancement. Using this integrated strategy, we show that Hoti-derived NP and Gc contribute distinct but complementary immune responses. To our knowledge, this study provides the first direct preclinical comparison of unformulated mRNA, DNA delivered by electroporation, and LNP-formulated mRNA for a CCHFV vaccine candidate.

## RESULTS

### 1. Screening of chimeric CCHFV vaccine genes

Six DNA vaccine candidates encoding Hoti strain-derived nucleoprotein (NP), GP38, and the Gc ectodomain (Gc_ecto) were generated and expressed either individually or as defined multi-antigen open reading frames separated by P2A self-cleaving peptides, with or without an N-terminal IgE leader (IgL) sequence to promote antigen secretion (Figure 1A). Candidates were evaluated *in vivo* following intramuscular immunization of C57BL/6J mice with 50 µg plasmid DNA delivered by electroporation at weeks 0 and 3 (Figure 1B). Following two immunizations, all NP-containing constructs induced high NP-specific IgG titers, whereas constructs encoding GP38 and/or Gc_ecto alone elicited comparatively lower antibody responses (Figure 1C). Inclusion of the IgL sequence enhanced antibody responses to secreted antigens, with the most pronounced effect observed for Gc_ecto-containing constructs (VIVAC-003.3 versus VIVAC-003.4; VIVAC-003.5 versus VIVAC-003.6). As expected, constructs lacking Gc_ecto (VIVAC-003.1 and VIVAC-003.2) did not induce detectable Gc-specific antibodies, confirming antigen specificity. GP38-specific antibodies were not detected in any group (Supplementary Figure S5). Cellular immune responses were assessed by IFN-γ ELISpot using antigen-specific peptide pools. NP-containing constructs elicited strong and reproducible NP-specific T cell responses across animals (Figure 1D), indicating efficient priming of cellular immunity. In contrast, GP38 did not induce detectable T cell responses (Supplementary Figure S6), and Gc_ecto alone contributed minimally to IFN-γ production, consistent with its primary role as a target of humoral rather than cellular immunity. Notably, constructs co-expressing NP and Gc_ecto elicited a complementary immune profile, integrating robust NP-driven T cell responses with enhanced Gc-directed antibody responses. Individual datasets for each immunization group are provided in Supplementary Figures S2-S6. Candidate prioritization was based on the combined magnitude and consistency of antigen-specific humoral and cellular responses across independent experiments. Among all constructs tested, VIVAC-003.3 exhibited the most balanced immune profile, characterized by strong NP-specific T cell responses together with high Gc-specific IgG titers (Figure 1C and 1D). Given that Gc is a surface-exposed envelope glycoprotein generated by glycoprotein precursor cleavage and mediates membrane fusion during viral entry, this immunological profile supports its relevance as a target of neutralizing antibodies. Based on this integrated immunogenicity profile, VIVAC-003.3 was selected for subsequent formulation and evaluation as an mRNA-LNP vaccine.

**Figure 1.**
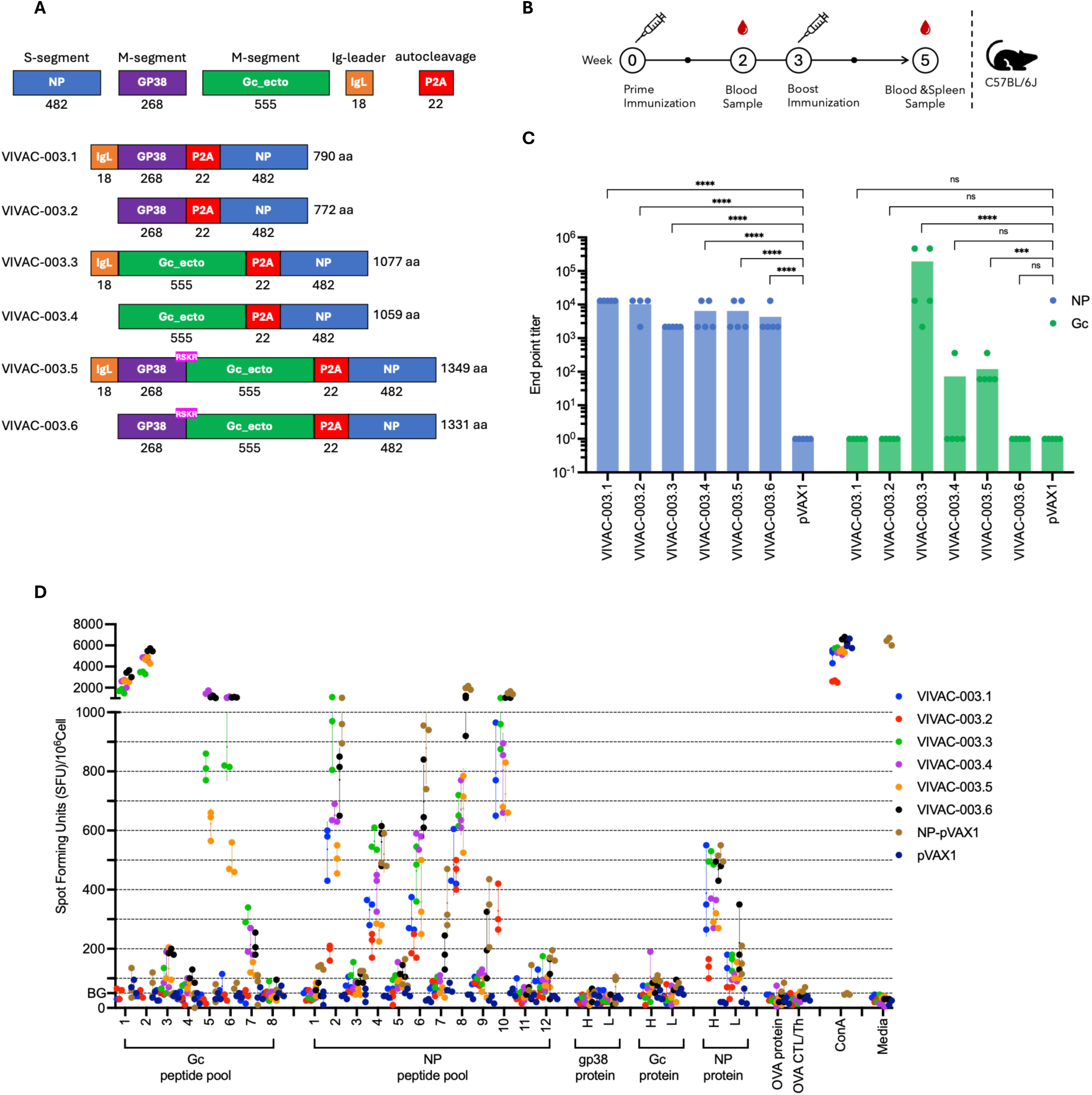
Vaccine candidate screening. **(A)** Hoti strain–derived antigens nucleoprotein (NP), GP38, and the Gc ectodomain (Gc_ecto) were expressed individually or in defined combinations as single open reading frames linked by P2A self-cleaving peptides. Constructs encoded GP38–P2A–NP, Gc_ecto–P2A–NP, or GP38–P2A–Gc_ecto–P2A–NP, each generated with or without an N-terminal IgG leader sequence (IgL) to modulate antigen secretion. All constructs were codon-optimized for expression in human cells. **(B)** *In vivo* immunization study design. C57BL/6J (H-2ᵇ) mice (n = 5 per group) were assigned to seven groups and immunized intramuscularly by *in vivo* electroporation at weeks 0 and 3 with six vaccine candidates or a pVAX1 vector control. Blood samples were collected at weeks 2 and 5, and spleens were harvested at study termination (week 5). **(C)** Antigen-specific antibody endpoint IgG titers against the indicated CCHFV antigens were measured by ELISA in sera collected at week 5 after two immunizations. Bars represent group means, with individual data points shown for each mouse (n = 5 per group). Endpoint titers were defined as the highest serum dilution exceeding the assay cutoff. Statistical significance was determined by one-way ANOVA with Dunnett’s multiple comparisons test (*p < 0.05; **p < 0.01; ***p < 0.001; ****p < 0.0001; ns, not significant). **(D)** IFN-γ–secreting cells were quantified by ELISpot and reported as spot-forming units (SFU) per 10⁶ splenocytes. Responses exceeding the mean of the medium-only background (BG) control were considered positive. Splenocytes were stimulated with Gc peptide pool (80 μg/ml), NP peptide pool (40 μg/ml), GP38 protein (1 μg/ml high, 0.1 μg/ml low), Gc protein (1 μg/ml high, 0.1 μg/ml low), N protein (1 μg/ml high, 0.1 μg/ml low), OVA protein (10 μg/ml), OVA CTL/Th peptides (10 μg/ml), or Concanavalin-A (ConA, 2 μg/ml). All stimulations were performed in triplicate, simble bars show mean and SD (n = 3).

### 2. Immunogenicity Comparison of Unformulated mRNA, Electroporated DNA, and mRNA-LNP

The selected VIVAC-003.3 antigen was produced as an mRNA-LNP vaccine using an in-house integrated manufacturing and analytical pipeline (Figure 2A). DNA-optimized VIVAC-003.3 (Figure 2B) was generated by IVT incorporating N1-methylpseudouridine, 3ʹ-O-methylated Cap 1 chemistry, and a template-encoded poly(A) tail, followed by oligo(dT) affinity purification. Analytical characterization confirmed production of full-length VIVAC-003.3 mRNA with minimal residual dsRNA and protein impurities (Supplementary Figure S7). Multiple independent mRNA-LNP batches were subsequently generated by microfluidic formulation, demonstrating consistent nanoparticle characteristics, with particle diameters within the expected range (67.7 ± 0.34 nm), low polydispersity (PDI 0.24 ± 0.01), and high encapsulation efficiency (>95%) (Supplementary Figure S7). These results demonstrate the reproducibility of the manufacturing workflow and support the suitability of the VIVAC-003.3 mRNA-LNP formulation for in vivo evaluation. C57BL/6J mice were immunized intramuscularly at weeks 0 and 3 with 2 µg of mRNA-LNP, 2 µg of unformulated mRNA, or 50 µg of plasmid DNA delivered by in vivo electroporation (Figure 2C). Sera and splenocytes were collected at week 5 to assess antigen-specific antibody and T cell responses. Humoral responses were measured by ELISA (Figure 2D). Mice immunized with mRNA-LNP developed Gc- and NP-specific IgG titers comparable to those induced by DNA vaccination, despite receiving a substantially lower vaccine dose (2 µg mRNA-LNP versus 50 µg DNA). Unformulated mRNA induced negligible responses, whereas DNA elicited detectable NP-specific antibody responses by week 2, which were not observed at this time point following mRNA-LNP immunization. By week 5, both platforms induced NP- and Gc-specific antibodies, with lower Gc-directed titers observed in the DNA group. Cellular immunity was quantified by IFN-γ ELISpot using NP and Gc_ecto peptide pools (Figure 2E), which demonstrated robust NP-specific T cell responses across groups, whereas Gc peptides contributed minimally overall. Notably, Gc pool 3 elicited exceptionally high responses, likely because it contains LSQMYSPVFEYLSGDR, which encompasses the conserved immunodominant CD8⁺ T-cell epitope YSPVFEYL previously mapped by Rao et al ^16^. Gc peptide pool 9 also elicited a strong IFN-γ response in C57BL/6 mice, whereas no comparable response was detected in BALB/c mice in the comparative study (Supplementary Figure S12A), consistent with the strain-dependent recognition. NP-driven responses were reproducible across animals. Individual ELISA and ELISpot data for each group are provided in Supplementary Figures S9-S11. A dose-matched comparison study performed in both C57BL/6 (H-2^b^) and BALB/c (H-2^d^) mice showed a similar overall immunogenicity profile. Comparative immunogenicity analysis revealed broadly similar response patterns in BALB/c and C57BL/6J mice, although the magnitude and antigen specificity differed between strains. BALB/c mice mounted higher Gc-specific antibody titers, whereas C57BL/6J mice showed stronger NP-specific antibody responses and generally more pronounced IFN-γ-secreting T-cell responses (Supplementary Figure S12). To assess whether the absence of immune responses following unformulated mRNA administration reflected a lack of *in vivo* translation, protein expression was evaluated using a firefly luciferase mRNA reporter. *In vivo* imaging demonstrated detectable luciferase expression at the injection site following administration of unformulated mRNA (Supplementary Figure S13). Despite measurable protein expression, unformulated mRNA failed to elicit antigen-specific humoral or cellular immune responses (Figure 2D and 2E). Functional neutralization was evaluated using a CCHFV tc-VLP assay (Figure 2F). Sera from mice immunized with mRNA-LNP strongly inhibited CCHFV tc-VLP entry, with neutralizing activity approaching that of the human positive-control serum, whereas sera from the DNA/E.P. group showed modestly lower activity. NT50 values could not be determined because the neutralization curves did not reach a defined lower plateau within the tested dilution range. In contrast, unformulated mRNA induced no detectable neutralizing activity, consistent with naïve mouse sera and the human negative-control serum. Overall, these findings demonstrate that both DNA and LNP-formulated VIVAC-003.3 mRNA elicit robust NP-specific T cell responses and potent Gc-directed antibody responses, resulting in a complementary immune profile with effective neutralizing activity against CCHFV.

**Figure 2.**
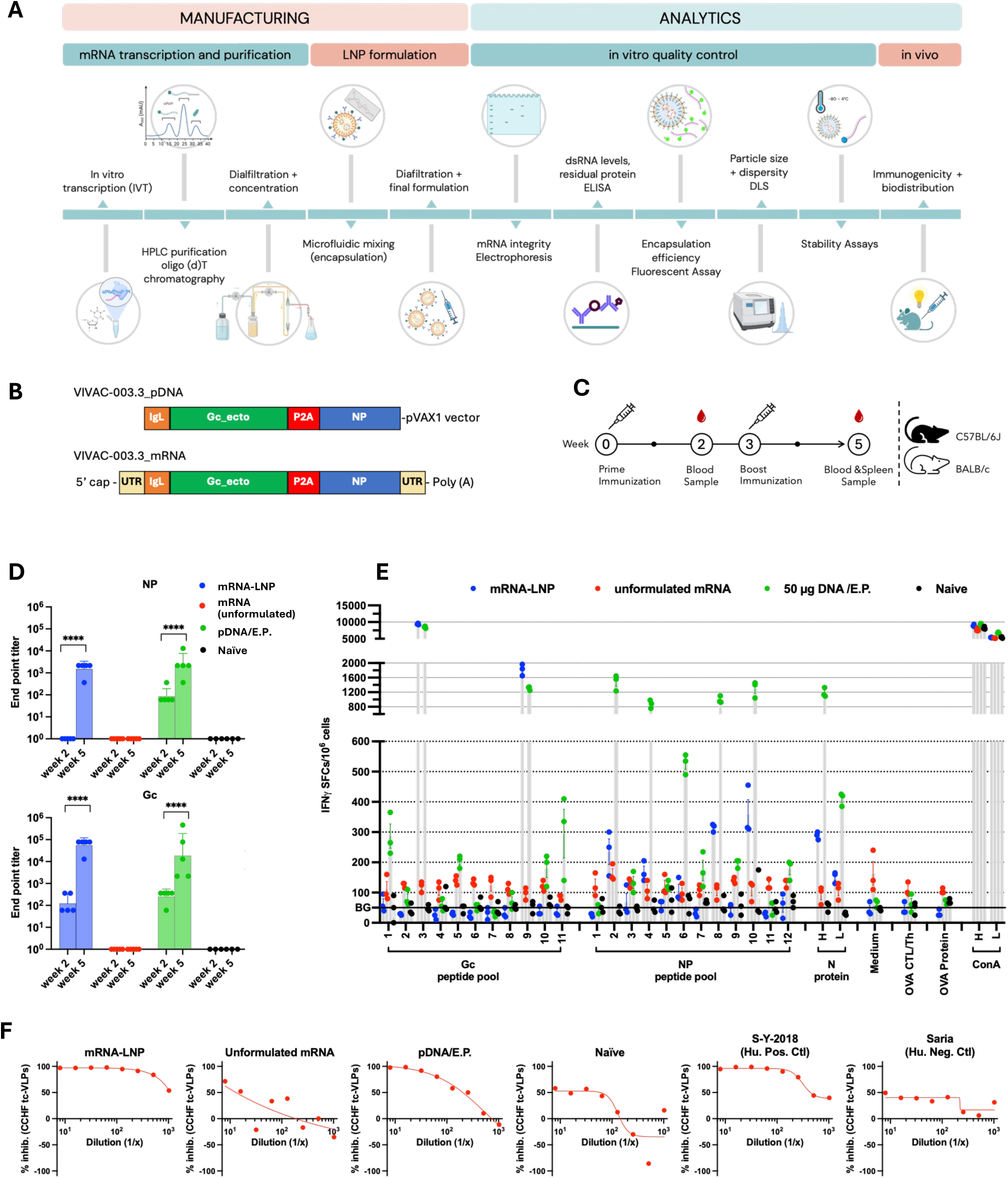
Comparative immunogenicity of DNA, unformulated mRNA, and LNP-formulated mRNA vaccines *in vivo*. **(A)** Overview of the mRNA–LNP manufacturing and analytical workflow. Manufacturing begins with in vitro transcription (IVT) of mRNA, followed by HPLC purification to remove impurities. The purified mRNA is then diafiltrated and concentrated, followed by microfluidic mixing for lipid nanoparticle (LNP) encapsulation and a final diafiltration step. Figure created with BioRender. **(B)** Schematic representation of DNA and mRNA vaccine constructs used for platform comparison. Plasmid DNA (pDNA) vaccines were administered with *in vivo* electroporation, while mRNA was evaluated either unformulated or formulated into lipid nanoparticles (LNPs). **(C)** *In vivo* immunization study design. C57BL/6J (H-2ᵇ) mice (n = 5 per group) were assigned to four groups and immunized intramuscularly at weeks 0 and 3 with pDNA (with electroporation), unformulated mRNA, mRNA-LNP, or left naïve as controls. Blood samples were collected at weeks 2 and 5, and spleens were harvested at study termination (week 5). **(D)** Antigen-specific serum IgG endpoint titers measured by ELISA at weeks 2 and 5 following immunization. Bars represent group means with individual mouse data points shown (n = 5 per group). Endpoint titers were defined as the highest serum dilution exceeding the assay cutoff. Statistical significance was determined by one-way ANOVA with Dunnett’s multiple comparisons test (*p < 0.05; **p < 0.01; ***p < 0.001; ****p < 0.0001; ns, not significant). **(E)** Quantification of antigen-specific IFN-γ–secreting splenocytes by ELISpot, reported as spot-forming units (SFU) per 10⁶ splenocytes. Responses exceeding the medium-only background (BG) control were considered positive. Splenocytes were stimulated with Gc peptide pool (80 μg/ml), NP peptide pool (40 μg/ml), N protein (1 μg/ml high, 0.1 μg/ml low), OVA protein (10 μg/ml), OVA CTL/Th peptides (10 μg/ml), or Concanavalin-A (ConA; 2 μg/ml high, 0.5 μg/ml low). All stimulations were performed in triplicate, symbol bars show mean and SD (n = 3). **(F)** Neutralizing activity of sera from vaccinated mice measured by a CCHF tc-VLP neutralization assay. Serial dilutions of pooled sera from mice immunized with mRNA–LNP, unformulated mRNA, or DNA/E.P. were incubated with CCHF transcriptionally-competent virus-like particles (tc-VLPs) before infection of target cells. Neutralization was quantified by luciferase reporter activity and the percentage of inhibition was calculated compared to untreated tc-VLPs. Human positive and negative control sera, and naive mouse sera were included as assay controls. Curves were fitted by nonlinear regression (4 parameters).

### 3. Protection against lethal homologous and heterologous CCHFV challenge

To evaluate whether the VIVAC-003.3 mRNA-LNP vaccine confers protection against lethal CCHFV infection, A129 IFNAR^−/−^ mice were immunized and subsequently challenged with two genetically distinct CCHFV strains. Mice received two intramuscular immunizations, administered three weeks apart, with either 2 µg or 6 µg VIVAC-003.3 LNP-mRNA. Negative-control animals received 6 µg Fluc LNP-mRNA. Two weeks after the booster immunization, mice were subjected to lethal viral challenge (Figure 3A and 3B). Groups 1-3 were challenged with the Hoti strain, whereas groups 4-7 were challenged with the IbAr strain. Group 7 served as a positive-control vaccine group and received a combination of 10 µg M and 10 µg NP mRNA vaccines based on the IbAr10200 strain (Figure 3A), a regimen previously shown to confer complete protection against CCHFV challenge ^19^. To determine whether immunization elicited antigen-specific humoral responses, antibodies against CCHFV Gc and N were measured by ELISA on the serum collected before challenge (Figure 3C-3F). Gc-specific antibody responses were detectable after a single immunization (T1, Figure 3C) in both the 2 µg and 6 µg VIVAC-003.3 LNP-mRNA groups and increased further after the second immunization (T2, Figure 3E). The 6 µg dose generally induced slightly higher Gc-specific responses than the 2 µg dose. However, group 5 showed unexpectedly lower Gc levels than group 2, despite these groups being designed to generate comparable responses. Gc-specific antibody levels appeared to associate with the subsequent challenge outcome. One animal in group 4 showed approximately 10% body-weight loss (Figure 3G) after challenge but recovered, coinciding with a high pre-challenge Gc antibody titer (Figure 3E). In contrast, animal 5_2 in group 5, which had the lowest Gc-specific antibody within that group, reached the humane endpoint (Figure 3G and 3E). All other challenged animals survived (Figure 3H). N-specific antibody responses remained low after the first immunization (Figure 3D) but increased in selected animals after the booster immunization, particularly in the 6 µg groups (Figure 3F). The positive-control group generated higher antigen-specific responses, consistent with vaccination using separate 10 µg M and 10 µg N DNA vaccine components. As expected, negative-control animals did not develop detectable CCHFV-specific antibodies and reached humane endpoints by day 4 after challenge (Figure 3H). These data demonstrate that 6 µg of mRNA-LNP delivered Gc-NP fusion antigen induces protective immunity effective against lethal challenge with two distinct CCHFV strains.

**Figure 3.**
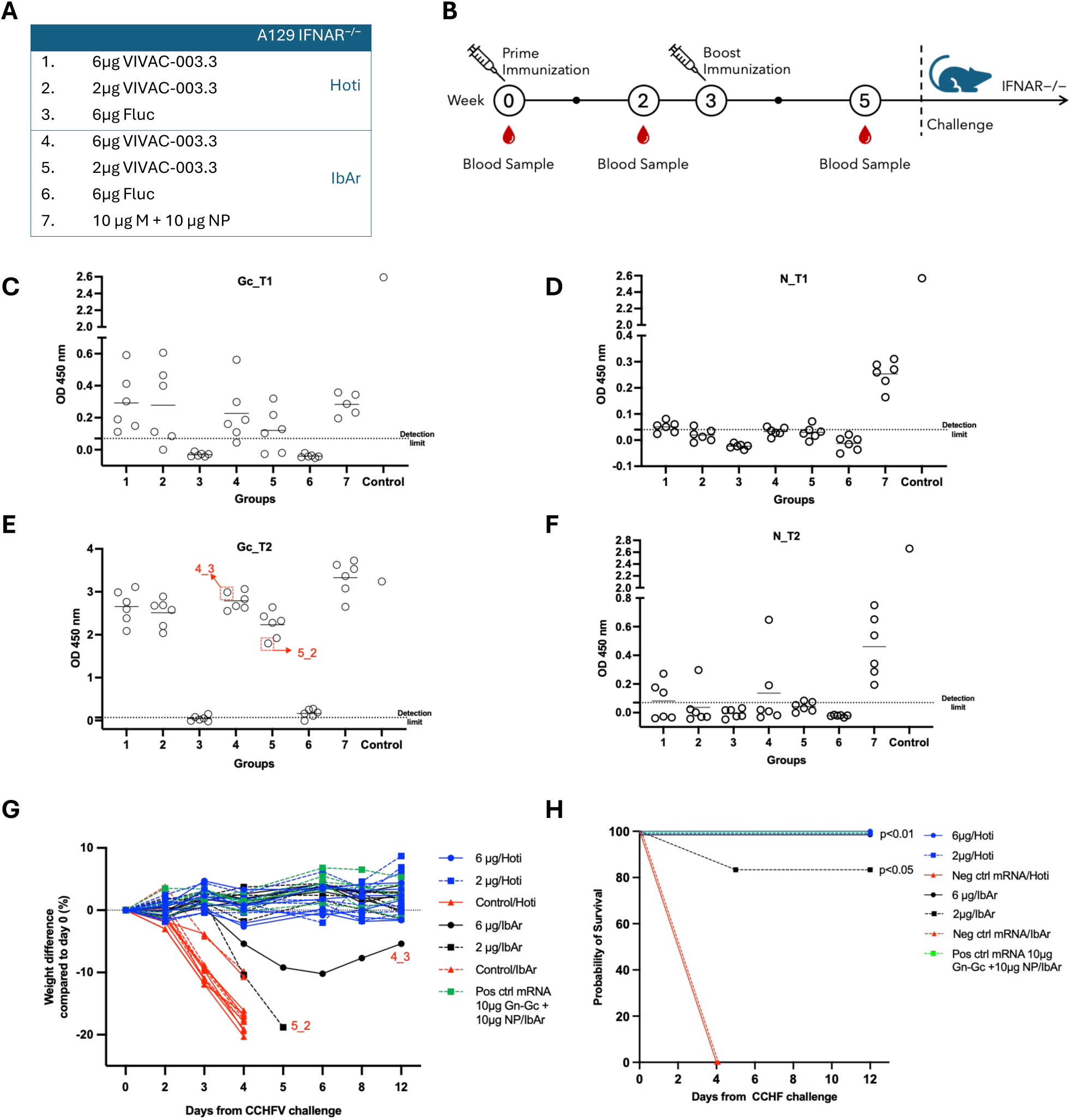
Immunogenicity and protective efficacy of the SVF-003.3 mRNA–LNP vaccine against CCHFV challenge. **(A)** Experimental groups of A129 IFNAR-/- mice immunized with 2 or 6 µg SVF-003.3 mRNA–LNP, control Fluc mRNA–LNP, or positive-control mRNAs encoding full M genome and the full N genome. Groups 1–3 were challenged with the CCHFV Hoti strain, whereas groups 4–7 were challenged with the IbAr10200 strain. **(B)** Immunization, blood-sampling, and challenge schedule. Mice received two immunizations, and challenge was performed two weeks after the second immunization. **(C–F)** Serum Gc-specific (C and E) and NP-specific (D and F) IgG responses after the first (T1; C and D) and second (T2; E and F) immunizations were measured by ELISA at a serum dilution of 1:100 and expressed as OD450 values. Each symbol represents an individual animal, and horizontal bars indicate group means. **(G)** Percentage change in body weight following CCHFV challenge. Individual animals are shown, with animals reaching the humane endpoint indicated by termination of the corresponding curves. **(H)** Kaplan–Meier survival curves during the 12-day observation period. Statistical significance relative to the corresponding Fluc control group is indicated in the graph.

## DISCUSSION

The nucleoprotein encoded by the S gene and the envelope glycoproteins encoded by the M segment are the primary antigens explored in CCHFV vaccine development. NP and M-segment-encoded glycoproteins have been evaluated, individually or in combination, across DNA, RNA, viral-vectored, and protein-subunit vaccine platforms ^1,20,26–28^. The surface-exposed glycoproteins Gn and Gc mediate viral attachment and membrane fusion, with Gc representing the best-established target of neutralizing antibodies ^12,17,18^. In contrast, NP is highly conserved across CCHFV phylogenetic clades and can induce strong antigen-specific cellular and non-neutralizing antibody responses ^13,14,24,28^. This conservation contrasts with the greater sequence variability of M-segment-encoded glycoproteins and may favour broader cross-strain immune recognition. Nevertheless, the magnitude and protective contribution of glycoprotein-directed responses vary according to antigen design and delivery platform ^24,27,29^. Preclinical studies have shown that Hoti-derived NP can confer protection against heterologous CCHFV challenge when delivered using replicating RNA, VSV-vectored, and recombinant protein vaccine platforms ^15,24,27^. In the recombinant subunit setting, NP alone prevented lethal disease, whereas the addition of GP38 further reduced morbidity among surviving animals ^24^. Collectively, these findings support the inclusion of conserved NP together with glycoprotein antigens in multicomponent vaccine designs. Recent RNA vaccine studies using either co-administered NP- and glycoprotein-encoding RNAs or single-RNA dual-antigen designs have induced coordinated humoral and cellular immune responses and conferred protection, reinforcing the rationale for combined antigen strategies ^19,29,30^. However, most previous approaches used separately encoded antigens ^15,26^ or full-length GPC ^25^, rather than a single fusion antigen combining NP with the Gc ectodomain ^19^. Although non-structural antigens such as GP38 have been proposed as supplementary targets, their contribution to immune protection appears variable, and GP38 did not induce robust immunity in the present antigen design and vaccination setting ^24,25,31^. Consistent with this literature, our findings demonstrate that Hoti-derived NP and the Gc ectodomain contribute distinct and complementary immunological functions, with NP driving robust antigen-specific T cell responses and Gc inducing strong neutralizing antibody responses, whereas GP38 did not elicit robust immunity in the present vaccine setting. In addition, incorporation of both antigens into a single fusion construct may streamline manufacturing, formulation, and quality control compared with the production and co-delivery of separate antigen constructs. These practical advantages, together with the observed immunogenicity and protective efficacy, support VIVAC-003.3 as a promising candidate for further preclinical development.

DNA and mRNA vaccines share the advantage that, unlike many viral-vector platforms, they can generally be readministered without the constraints imposed by classical anti-vector immunity ^32^. Although they both rely on *in vivo* antigen production, they differ significantly in several important aspects, including their delivery methods, dosage needs, immunogenic profiles, duration of immune response, safety profiles, and manufacturing processes ^32^. Regarding delivery, plasmid DNA needs to enter the nucleus to be transcribed. Electroporation is commonly used to greatly enhance cellular uptake and subsequent antigen expression ^33,34^. However, electroporation requires specialized devices, trained personnel, and standardized administration conditions, adding operational complexity that may affect large-scale deployment, particularly in resource-limited settings ^35,36^. In contrast, mRNA vaccines, typically encapsulated in LNPs, are delivered directly to the cytosol. Following endosomal escape, the mRNA is translated, and these formulations can be administered via standard injection ^37,38^. Effective dose requirements cannot be generalized solely according to platform, because they depend on antigen design, delivery route, formulation, and expression efficiency. However, mRNA-LNP vaccines frequently induce strong immune responses at lower nominal nucleic acid doses than plasmid DNA vaccines (Figure 2) ^32,39^. A head-to-head SARS-CoV-2 study across hamster, guinea pig, rabbit, and mouse models reported stronger cellular responses following DNA-EP vaccination and more pronounced humoral responses following mRNA vaccination, although differences in dose, formulation, and administration route preclude attribution of these effects exclusively to the vaccine platform ^39^. Consistent with these observations, in a comparison using a separate vaccine candidate, we evaluated 2 µg plasmid DNA delivered by electroporation versus 2 µg mRNA-LNP. While a 2 µg DNA dose was sufficient to elicit detectable immune responses, the magnitude of the response was lower than that induced by 2 µg mRNA-LNP, indicating greater immunogenicity of the mRNA-LNP formulation at the tested mass dose (Figure S14). Regarding safety, DNA vaccination delivered by electroporation is generally well tolerated but is associated with transient local tissue effects and requires controlled device use ^35–37^. Since the widespread deployment of mRNA vaccines during the COVID-19 pandemic, extensive mechanistic, clinical, and post-authorisation evidence has established mRNA vaccination as a clinically validated and adaptable platform with a generally favourable benefit-risk profile ^40,41^. Nevertheless, mRNA is not intrinsically superior for every antigen or disease, and the safety and performance of licensed products cannot be extrapolated directly to all future mRNA constructs. In particular, the biological activity of mRNA-LNP vaccines is formulation dependent. Empty LNPs can activate and promote the maturation of antigen-presenting cells ^42^, whereas certain LNP compositions have produced pronounced inflammatory responses in preclinical models ^43^. Clinical experience has also identified rare hypersensitivity reactions associated with PEG-containing formulations, with anti-PEG antibodies proposed as one potential contributing mechanism ^44^. Accordingly, each mRNA vaccine should be evaluated as an individual product, considering the encoded sequence, RNA integrity and impurities, LNP composition, biodistribution, dose, reactogenicity, immunogenicity, and protective efficacy ^41,45^. The same product-specific principle also applies to DNA vaccines, for which antigen design, plasmid backbone, manufacturing quality, dose, delivery route, and electroporation parameters can substantially influence both safety and immunogenicity ^32,35,46^. Thus, comparisons between DNA and mRNA vaccination should focus on the properties of the complete vaccine products rather than on platform classification alone. In terms of durability, DNA vaccines delivered by electroporation have demonstrated sustained cellular immune responses in both preclinical models and clinical studies ^34,47,48^. For licensed SARS-CoV-2 mRNA vaccines, strong peak antibody responses are followed by a measurable decline in circulating antibody titers, although durability varies substantially according to product, dose, immunization schedule, prior infection, and host factors ^49,50^. Accordingly, immunogenicity profiles differ but overlap, as DNA-electroporation strategies often favor robust T cell responses ^33,34,51^, whereas mRNA-LNP platforms frequently elicit robust humoral responses ^37,38^. These tendencies are not absolute and can be modified substantially by antigen composition, formulation, dose, and delivery route. A direct preclinical comparison similarly found stronger T-cell responses following DNA vaccination and stronger antibody responses following mRNA vaccination ^39^. Finally, from a manufacturing perspective, DNA vaccines are produced via bacterial fermentation with established purification and quality-control frameworks ^46,52,53^, In contrast, mRNA vaccines are produced using a cell-free, enzymatic in vitro transcription process that enables rapid sequence modification and greater adaptability ^45,54^. Taken together, these considerations provide a framework for interpreting the performance of the DNA-electroporation and mRNA-LNP formulations evaluated in this study. The relative suitability of the two approaches will also depend on the balance among immunogenicity profile, effective dose, durability, safety, delivery requirements, and manufacturing considerations in the intended setting.

To support the translational development of the selected candidate we established a scalable mRNA-LNP manufacturing and analytical workflow designed to support progression from research-scale candidate development toward clinical translation and future GMP manufacturing within the Karolinska Advanced Therapy Medicinal Products (ATMP) Center, (Figure 2A). This platform provides an integrated framework for mRNA production, LNP formulation, and analytical characterization, enabling the rapid production and evaluation of vaccine candidates such as VIVAC-003.3. The development of scalable manufacturing processes and associated analytical methods is an important requirement for advancing mRNA products toward early-phase clinical evaluation ^54^ and aligns with current priorities for accelerating CCHFV vaccine development. A detailed methodological description of the platform will be reported separately.

However, several limitations of this study should be acknowledged. First, immunogenicity and protective efficacy were evaluated in murine models, which provide valuable platforms for candidate screening but do not fully recapitulate the heterogeneity of human CCHF or predict clinical protection. Second, we did not directly assess local or systemic inflammatory cytokine profiles following DNA-EP versus mRNA-LNP vaccination, which could provide further insight into platform-specific innate immune activation and its relationship to subsequent adaptive immunity.

In summary, the Hoti-derived NP-Gc vaccine induced complementary immune responses, with NP mainly promoting T-cell immunity and Gc inducing neutralizing antibodies. The mRNA-LNP formulation produced strong immune responses at a lower dose and fully protected mice against both Hoti and IbAr CCHFV challenge. These findings support VIVAC-003.3 as a promising candidate for further preclinical development.

## METHODS AND MATERIALS

### DNA Constructs Design and Plasmid Preparation

Vaccine constructs were designed using antigens derived from the Hoti strain, including GP38, the Gc ectodomain, and NP. Antigens were assembled as single open reading frames linked by P2A self-cleaving peptides to enable coordinated expression. For selected constructs, an N-terminal IgE signal peptide was included to promote antigen secretion, whereas corresponding constructs lacking a signal peptide were generated for comparison. Specifically, constructs encoded GP38-P2A-NP or Gc ectodomain-P2A-NP, with or without an N-terminal IgE signal sequence. In addition, a trivalent construct encoding GP38-P2A-Gc ectodomain-P2A-NP was generated, again in versions containing or lacking the IgL (Figure 1A). All sequences were codon-optimized for human cell expression and synthesized as plasmid DNA (GenScript). Plasmids were propagated in *Escherichia coli* TOP10 cells and purified using endotoxin-free purification reagents (Qiagen) according to the manufacturer’s instructions. The expected restriction pattern of the construct was confirmed by diagnostic digestion with BamHI and XbaI. (FastDigest, Thermo Fisher Scientific).

### mRNA Design, Synthesis, and Purification

CCHF mRNA was synthesized by *in vitro* transcription (IVT) from the linearized double-stranded DNA plasmid encoding the CCHF gene plasmid (GenScript, VIVAC-003.3 U9360GPAG0) consisting of a T7 promoter followed by a synthetic 5ʹ UTR containing a Kozak translation initiation sequence, the optimized coding sequence, a β-globin-derived 3ʹ UTR, and a 100-nucleotide poly(A) tail (GenScript). Modified mRNA was transcribed using the CleanScript™ IVT Kit (TriLink, K-7413) with N1-methyl-pseudouridine (m1Ψ), a 3ʹOMe-CleanCap, 100 ng/μL linearised template DNA, and incubated at 37°C for 3 h with 300 rpm horizontal shaking. Following transcription, the DNA template was digested with DNase I-XT (New England Biolabs) at 37 °C for 15 minutes, after which the reaction was terminated with 50 mM EDTA and 500 mM NaCl. The crude mRNA was purified by affinity HPLC on a POROS GoPure Oligo(dT)₂₅ column (5 mL; Thermo Fisher Scientific) using an ÄKTA Pure 25 chromatography system. The RNA was equilibrated and loaded in Buffer A (10 mM Tris-HCl, 1 mM EDTA, 500 mM NaCl, pH 7.4), washed with Buffer B (10 mM Tris-HCl, 1 mM EDTA, pH 7.4), and the RNA was eluted isocratically in nuclease-free water. Eluted fractions were monitored by UV absorption at 260 nm, and peak fractions were analyzed by agarose gel electrophoresis to confirm RNA integrity and purity. Fractions containing intact, pure mRNA were pooled, diafiltrated into H_2_O, concentrated using Amicon Ultra centrifugal filters with a 100 kDa molecular weight cut-off (Merck Millipore), and frozen in aliquots at −20 °C until use.

### LNP Formulation

Purified mRNA was encapsulated in LNPs using the NanoAssemblr™ Ignite™ system (Cytiva, NIN0001) equipped with NxGen™ microfluidic cartridges. The CCHF mRNA was formulated using the RNA Delivery LNP Kit (Cytiva; 1002471) at a nitrogen-to-phosphate (N:P) ratio of 8, whereas the OVA mRNA was formulated using GenVoy-ILM™ lipid mix (Precision NanoSystems/Cytiva; NWW0041) at an N:P ratio of 4. For both formulations, mRNA diluted in formulation buffer (Cytiva; 1002471) was mixed with the lipid solution prepared in ethanol at an aqueous-to-organic flow rate ratio (FRR) of 3:1 and a total flow rate (TFR) of 20 mL min⁻¹. Following formulation, LNPs were concentrated to approximately 400 µL using Amicon® Ultra centrifugal filters (100 kDa molecular weight cut-off; MilliporeSigma) and stored at 4 °C until use.

### mRNA-LNP Quality Control

mRNA quality was assessed prior to LNP formulation. RNA purity was evaluated by measuring absorbance ratios (A260/A280 and A260/A230) using a NanoDrop™ One spectrophotometer (Thermo Fisher Scientific). mRNA integrity was assessed by agarose gel electrophoresis (1% (w/v) agarose gel, 160 V for 30 min). Double-stranded RNA (dsRNA) impurities were quantified using the Lumit™ dsRNA Detection Kit (Promega) according to the manufacturer’s instructions, and residual protein content was assessed using the AccuOrange™ Protein Quantitation Assay.

Following LNP formulation, particle size (Z-average hydrodynamic diameter) and polydispersity index (PDI) were determined by dynamic light scattering (DLS) using a ZetaStar™ instrument (Waters Corporation). Encapsulation efficiency was determined by quantifying total and unencapsulated RNA using the Quant-iT™ RiboGreen™ RNA Assay Kit (Thermo Fisher Scientific; R11490). Total RNA was quantified following LNP disruption with 2% (v/v) Triton™ X-100 in TE buffer, whereas free RNA was quantified without detergent treatment. Encapsulation efficiency was calculated as [(total RNA − free RNA) / total RNA] × 100.

### Cell culture and transfection

HEK293T cells were seeded at 5×10^5^ cells per well in 12-well plates and cultured overnight at 37°C with 5% CO₂. Cells were transfected with 2 µg mRNA-LNP, 2 µg mRNA complexed with Lipofectamine™ MessengerMAX™ Transfection Reagent (Invitrogen™, LMRNA001), or 2 µg plasmid DNA complexed with Lipofectamine™ 3000 Transfection Reagent (Invitrogen™, L3000008). Untreated cells served as negative controls. Cells were harvested 24 h post-transfection.

### Western blot

Cell pellets were lysed in protease inhibitor-supplemented lysis buffer for 30 min at 4°C. Clarified lysates were quantified by BCA assay, and 50 µg total protein per sample was separated on 4-12% Bis-Tris gels (Invitrogen™, NP0323BOX) and transferred to PVDF membranes. Proteins were transferred to PVDF membranes using an iBlot Dry Blotting System and iBlot PVDF Mini Transfer Stacks (Invitrogen™, IB24002) with program P4 at 15 V for 7 min. Membranes were blocked for 1 h at room temperature in 1× Pierce Clear Milk Blocking Buffer (Thermo Scientific™, 37587). Separate membranes were incubated with pooled mouse sera containing antibodies against both Gc and NP (in-house), diluted 1:1000 in blocking buffer, overnight at 4°C. Membranes were washed three times for 10 min with PBS containing 0.5% Tween-20 and subsequently incubated for 1 h at room temperature with HRP-conjugated anti-mouse secondary antibody (Invitrogen™, 61-6520) diluted 1:2000 in blocking buffer. After three additional 10-min washes, protein bands were detected using Pierce™ ECL Western Blotting Substrate (Thermo Scientific™, 32209) and imaged with a ChemiDoc Imaging System. Exposure times ranged from 5 s to 2 min.

### Animals

For the immunogenicity studies, female C57BL/6 (H-2^b^) and BALB/c (H-2^d^) mice (8-12 weeks) were obtained from Charles River Laboratories (Sulzfeld, Germany) and housed under standard conditions at the Preclinical Laboratory (PKL), Karolinska University Hospital, Huddinge. For the challenge study, female IFNAR^−/−^ (B6.129S2-Ifnar1tm1Agt/Mmjax strain) mice, aged 11-13 weeks at the start of immunization, were obtained from the Jackson Laboratory (No. #010830, USA). Animal studies involving infectious agents were performed in the biosafety level 4 (BSL-4) laboratory at the Public Health Agency of Sweden. Animals were maintained under standard housing conditions and monitored daily throughout the study. All animal procedures were approved by the Stockholm Animal Research Ethics Committee (Dnr 13278-2019, 16975-2024) and Linköping Animal Research Ethics Committee (15671-2024) and conducted in accordance with institutional and national guidelines.

### Immunization study

Female mice (5 per group) were immunized twice at three-week intervals and sacrificed two weeks after the final immunization. DNA immunizations were performed as previously described ^55,56^. Briefly, 50 μg of plasmid DNA in 50 μL sterile PBS was injected intramuscularly into the tibialis anterior (TA) muscle using a 27G needle, followed by *in vivo* electroporation using the Cliniporator2 device (IGEA) with a two-needle electrode. Analgesia was administered prior to injection, and mice were maintained under isoflurane anesthesia. Electroporation consisted of a single 1-ms 600 V/cm pulse followed by a single 400-ms 60 V/cm pulse to enhance plasmid uptake. For mRNA immunizations, mice received 2 μg of mRNA-LNP or unformulated mRNA in 50 μL sterile PBS by intramuscular injection. Injection sites were disinfected prior to administration. Mice were weighed before and three days post-injection, and well-being was monitored daily for three days following each immunization. Spleen and blood were collected as previously described ^55^.

### Challenge study

Female IFNAR^−/−^ mice, 11-13 weeks of age, were assigned to seven groups (n = 6 per group) and immunized twice at a 3-week interval. All mRNA-LNP vaccine formulations were diluted in calcium- and magnesium-free PBS and administered by intramuscular injection in a total volume of 50 μL per mouse. Blood samples were collected before the first immunization, before the booster immunization, and before viral challenge. Two weeks after the second immunization, mice were challenged intraperitoneally with 400 focus-forming units (FFU) of CCHFV in a total volume of 100 μL. After challenge, animals were monitored daily for up to 12 days for survival, body weight, clinical signs of disease, and overall welfare according to approved humane endpoint criteria. At study termination or when humane endpoints were reached, blood was collected and processed for serum separation.

### *In vivo* bioluminescence imaging

BALB/c (H-2^d^) mice (8-12 weeks old) were injected intramuscularly (i.m.) into the tibialis anterior muscle with luciferase-encoding nucleic acid constructs. Five experimental groups were included: 8 µg LNP-formulated mRNA (n = 5), 8 µg unformulated mRNA (n = 3), 8 µg unformulated mRNA delivered by electroporation (EP; n = 2), 8 µg plasmid DNA delivered by EP (n = 5), and a naïve control group receiving no injection (n = 5). For E.P. groups, electroporation was performed immediately following intramuscular injection using established parameters. For bioluminescence imaging, D-luciferin (firefly luciferin, potassium salt; PerkinElmer, #122799) was prepared fresh at 15 mg/mL in calcium- and magnesium-free PBS, filter-sterilized through a 0.2 µm syringe filter, and protected from light until use. Mice received luciferin by intraperitoneal injection at 150 mg/kg body weight (10 µL/g) 10-15 min before imaging. Bioluminescence imaging was performed using an *in vivo* imaging system (IVIS) on days 1, 2, 5, 7, and 12 post-administrations. Luminescence signals at the injection site were quantified and reported as radiance (photons/sec/cm²/sr).

### Detection of IgG-specific antibodies

Blood was collected by submandibular venipuncture and allowed to clot for at least 30 min at 4°C. Serum was isolated by two sequential centrifugation steps at 6,000 rpm for 6 min and stored until analysis.

Antigen-specific serum IgG responses against CCHFV NP and Gc were measured by indirect ELISA. Ninety-six-well MaxiSorp plates (Nunc; Invitrogen) were coated overnight at 4°C with 1 μg/mL recombinant NP or Gc protein (Native Antigen, UK; Ref: REC31639, REC31696) diluted in 50 mM sodium carbonate buffer (pH 9.6). Plates were washed three times with 200 μL PBS containing Tween 20 (PBST) after each incubation step.

For detection of NP-specific antibodies, plates were blocked with 10% skimmed milk powder (Fisher Scientific, 70166) in PBS for 1 h at 37°C. Serum samples were initially diluted 1:60 and serially diluted across the plate, followed by incubation for 1 h at 37°C. Bound antibodies were detected using rabbit anti-mouse IgG conjugated to horseradish peroxidase (1:1,000; Invitrogen, 61-6520) for 1 h at 37°C. Plates were developed with 3,3ʹ,5,5ʹ-tetramethylbenzidine substrate (Sigma-Aldrich, T0440) for 12 min, and the reaction was stopped using TMB stop reagent (Sigma-Aldrich, S5814). Absorbance was measured at 450 nm.

For detection of Gc-specific antibodies, plates were blocked with PBS-A containing 1% bovine serum albumin and 2% goat serum for 1 h at 37°C. Serially diluted serum samples were incubated for 1 h at 37°C, followed by goat anti-mouse IgG γ-chain-specific alkaline phosphatase conjugate (1:1,000; Sigma-Aldrich, A1047) for 1 h at 37°C. Plates were developed in the dark for 30 min with 100 μL/well p-nitrophenyl phosphate substrate, prepared by dissolving one 5-mg tablet (Sigma-Aldrich, S0942) in 5 mL of 1 M diethanolamine buffer containing 0.5 mM MgCl₂. The reaction was stopped with 50 μL of 1 M NaOH, and absorbance was measured at 405 nm with background correction at 620 nm.

Serum from a nonimmunized mouse was included as a negative control. Endpoint titers were defined as the highest serum dilution producing an absorbance value at least threefold greater than that of the negative-control serum at the corresponding dilution.

### Detection of specific IFN-γ producing T cells

Pre-coated IFN-γ ELISpot plates (#3321-4APW, Mabtech) were washed with PBS and blocked for ≥2 h at 37°C with complete RPMI 1640 (GlutaMAX™, 10% FBS, 1% HEPES, 1% non-essential amino acids, 1% penicillin/streptomycin, 1×10⁻⁶% 2-mercaptoethanol). Spleens were harvested, and single-cell suspensions were prepared at 2×10⁶ cells/mL. Cells were pooled by study group and seeded at 2×10⁵ cells/well. Antigens, including peptides and proteins were added at indicated concentrations; culture medium served as a negative control. The Gc ectodomain and N protein were divided into 20-mer peptides with 10-aa overlaps. Peptides were pooled into five peptides per Gc pool (11 pools total) and four per N pool (12 pools total). Plates were incubated 36-48 h at 37°C with 5% CO₂. Biotinylated anti-IFN-γ antibody (R4-6A2-biotin, #3321-3, Mabtech) at 2 μg/mL in PBS/0.5% FBS was added for 2 h at room temperature, followed by 0.1% Streptavidin-ALP (#3310-10-1000, Mabtech) for 1 h. Spots were developed using BCIP/NBT-plus substrate (#3650-10, Mabtech) for 12 min and washed with tap water. Plates were dried and analyzed using Mabtech IRIS™ 2 and Apex™ software.

### Neutralization Assay of CCHF transcriptionally-competent virus-like particles (tc-VLPs)

CCHFV-neutralizing antibody responses were assessed using a microneutralization assay based on IbAr10200 transcriptionally-competent virus-like particles (tc-VLPs) ^57^ containing a nanoluciferase reporter gene ^58^. C57BL/6J mice were immunized twice with 2 µg LNP-formulated mRNA, 2 µg unformulated mRNA, or 50 µg DNA delivered by intramuscular electroporation. Naïve mice were included as controls. Sera were collected, pooled within each vaccination group, and heat-inactivated at 56°C for 30 min.

Twenty hours before the microneutralization assay, HuH-7 cells were reverse-transfected with 100 ng pV5_CAGGS_CCHFV-IbAr_L-CO (GenBank: PZ941510) and 30 ng pCAGGS_CCHFV-IbAr_N-CO[SD1.1] (GenBank: PZ941511) per well of a 96-well plate, using GeneJammer transfection reagent (Agilent). The heat-inactivated sera were serial diluted 2-fold in DMEM, mixed with CCHF IbAr10200 tc-VLPs to reach final dilutions 1:8 to 1:1024, and incubated at 37°C for 1 h to allow antibody-mediated neutralization. The serum-tc-VLP mixtures were then added in technical duplicate to the pre-transfected HuH-7 cells, previously washed one time with PBS. After 1 h of incubation at 37°C, complete medium with 10% FBS was added to the inoculum and the cells were further incubated for 20 hours at 37°C, 5% CO2. The next day, the cells were lysed with Passive Lysis Buffer (Promega), and luciferase activity was detected using the Nano-Glo® luciferase assay system (Promega) and the Centro XS luminometer (Berthold Tech). The percentage inhibition was calculated for each duplicate, using the formula 100*(1-(x-min)/(max-min)), in which “min” is the mean value of the Mock wells, and “max” is the mean value of the “tc-VLP only” wells. A non-linear regression analysis (4 parameters) was done in GraphPad Prism.

### Statistical analysis

Data were analyzed on GraphPad Prism Version 10.6.1(799) software.

## DATA AND CODE AVAILABILITY

All raw data will be available upon request.

## ACKNOWLEDGMENTS

This work was supported by the European Union’s Horizon 2020 programme (Grant No. 732732) and Horizon Europe programme (Grant No. 101137033). The study was also supported by the Swedish Research Council (M.S. and A.M.), the Swedish Cancer Society (M.S.), Karolinska Institutet (M.S. and L.F.), Stockholm County Council CIMED (M.S), ALF grants (M.S.), and private donations to M.S. and A.M. We acknowledge the Karolinska ATMP Center for its infrastructure, technical expertise, and support in establishing the mRNA-LNP manufacturing and analytical pipeline.

## AUTHOR CONTRIBUTIONS

Conceptualization: M.S., A.M., G.A., J.Y.

Data curation: J.Y., G.A., C.N., L.S.

Formal analysis: J.Y., C.N., L.S., S.A., M.M., S.D., P.B.

Funding acquisition: M.S., A.M.

Investigation: J.Y., G.A., C.N., S.A., L.S., M.M., S.D., P.B., E.O.

Methodology: G.A., J.Y., C.N., S.A., L.S., S.D,

Project administration: M.S., G.A.

Resources: BioRender

Software: GraphPad Prism Version 10.6.1(799)

Supervision: M.S., G.A.

Validation: G.A., C.N., S.A., S.D., L.S.

Writing - original draft: J.Y

Writing - review and editing: M.S., A.M., G.A., C.N., L.S., M.M., S.A., S.D., L.F.

## DECLARATION ABOUT THE USE OF GENERATIVE AI

The AI-assisted tool OpenAI ChatGPT (GPT-5 model family) was used to edit the language and improve the clarity of author-provided text, and the final version was reviewed and approved by the authors.

---

**Supplementary Figure S1.**
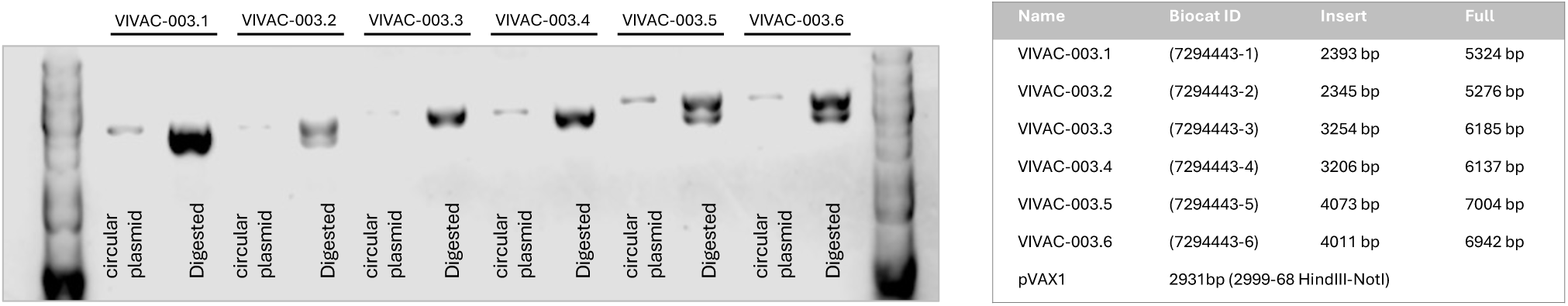
Agarose gel electrophoresis analysis of plasmid DNA constructs. This agarose gel showing the integrity of plasmid DNA constructs used in this study. For each construct, DNA was examined in both its circular and linearized forms. Undigested samples show the expected supercoiled and open circular conformations, while digested samples appear as single bands that match the predicted full-length plasmid sizes, as listed in the table below the gel. These results confirm the correct size and structural integrity of all plasmid preparations.

**Supplementary Figure S2.**
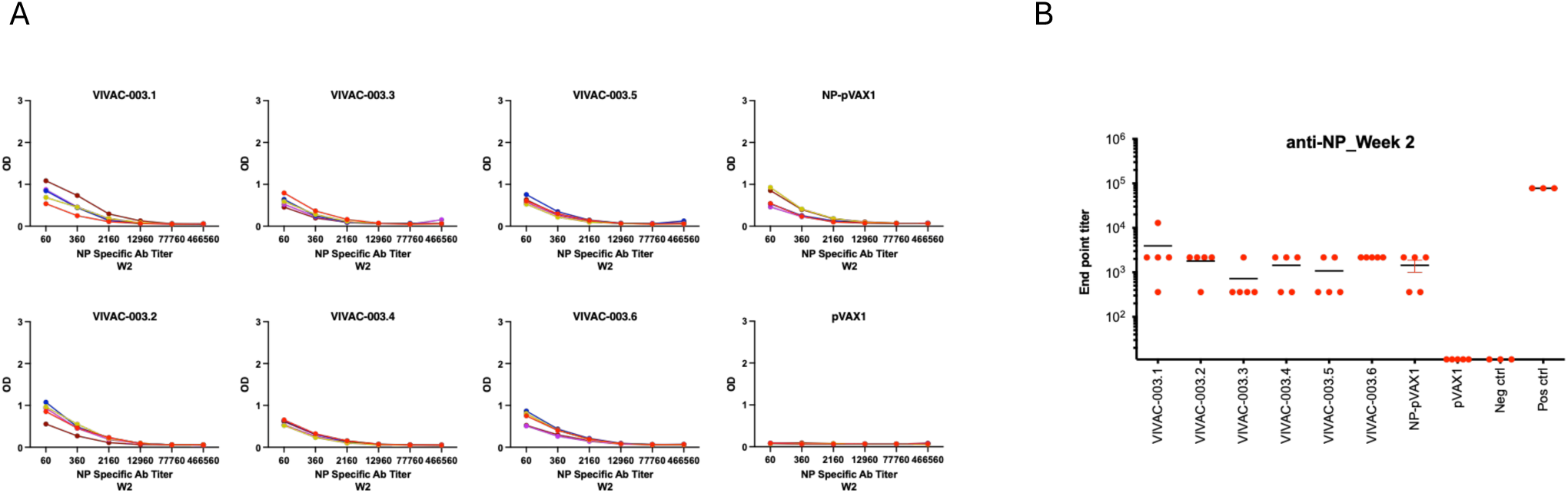
ELISA characterization of NP-specific IgG responses across vaccine constructs two weeks post vaccination. **A.** NP-specific serum IgG responses were quantified by ELISA using serially diluted sera collected from immunized mice. Optical density (OD) values were calculated as OD450–OD620 after background subtraction. **B.** Endpoint IgG titers for each group, with each dot representing an individual mouse and horizontal bars indicating group means (with SEM).

**Supplementary Figure S3.**
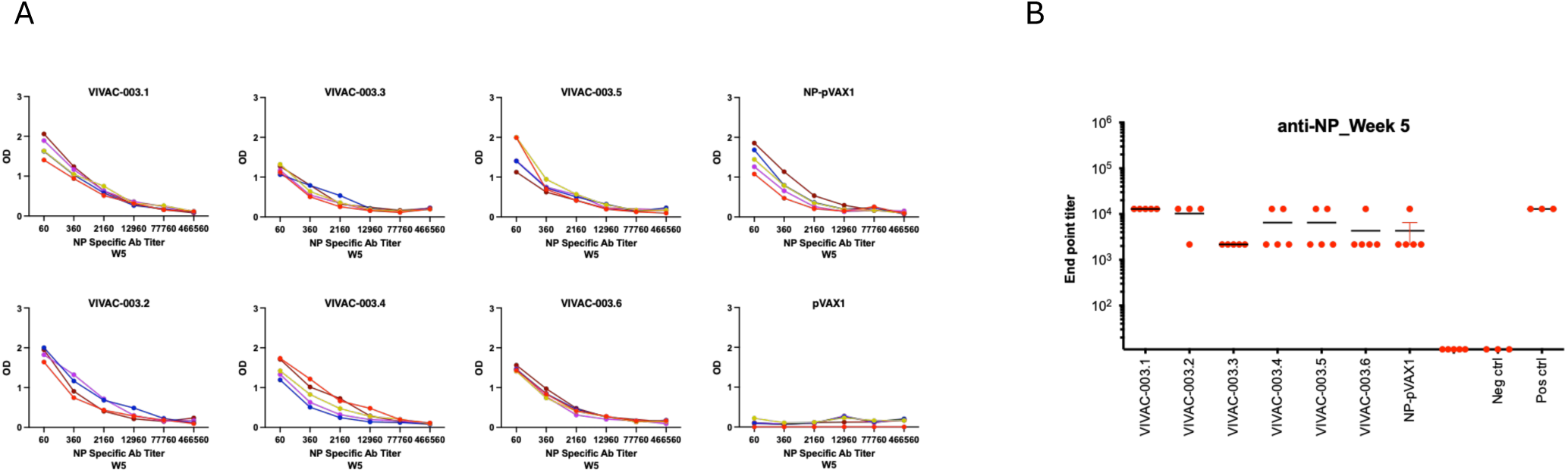
ELISA characterization of NP-specific IgG responses across vaccine constructs five weeks post vaccination. **A.** NP-specific serum IgG responses were quantified by ELISA using serially diluted sera collected from immunized mice. Optical density (OD) values were calculated as OD450–OD620 after background subtraction. **B.** Endpoint IgG titers for each group, with each dot representing an individual mouse and horizontal bars indicating group means (with SEM).

**Supplementary Figure S4.**
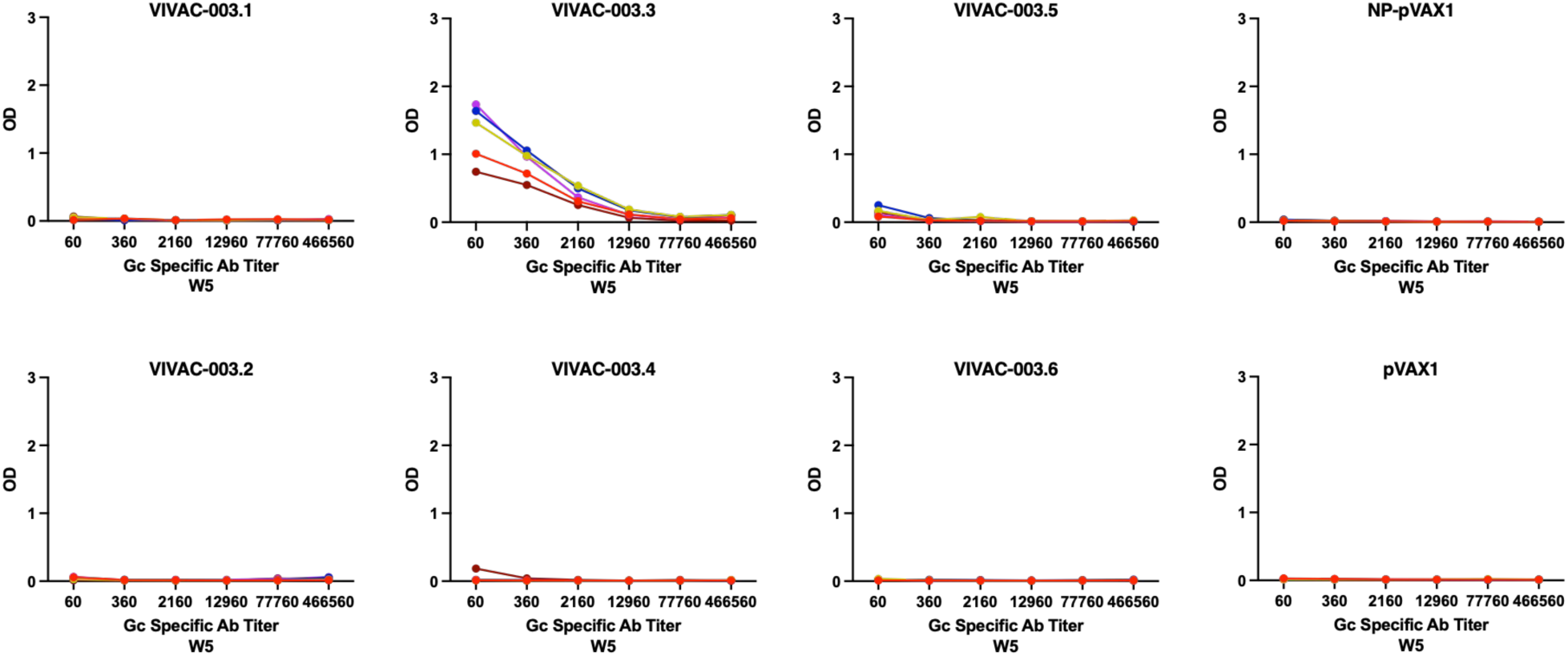
ELISA characterization of Gc-specific IgG responses across vaccine constructs five weeks post vaccination. Gc-specific serum IgG responses were quantified by ELISA using serially diluted sera collected from immunized mice. Optical density (OD) values were calculated as OD450–OD620 after background subtraction.

**Supplementary Figure S5.**
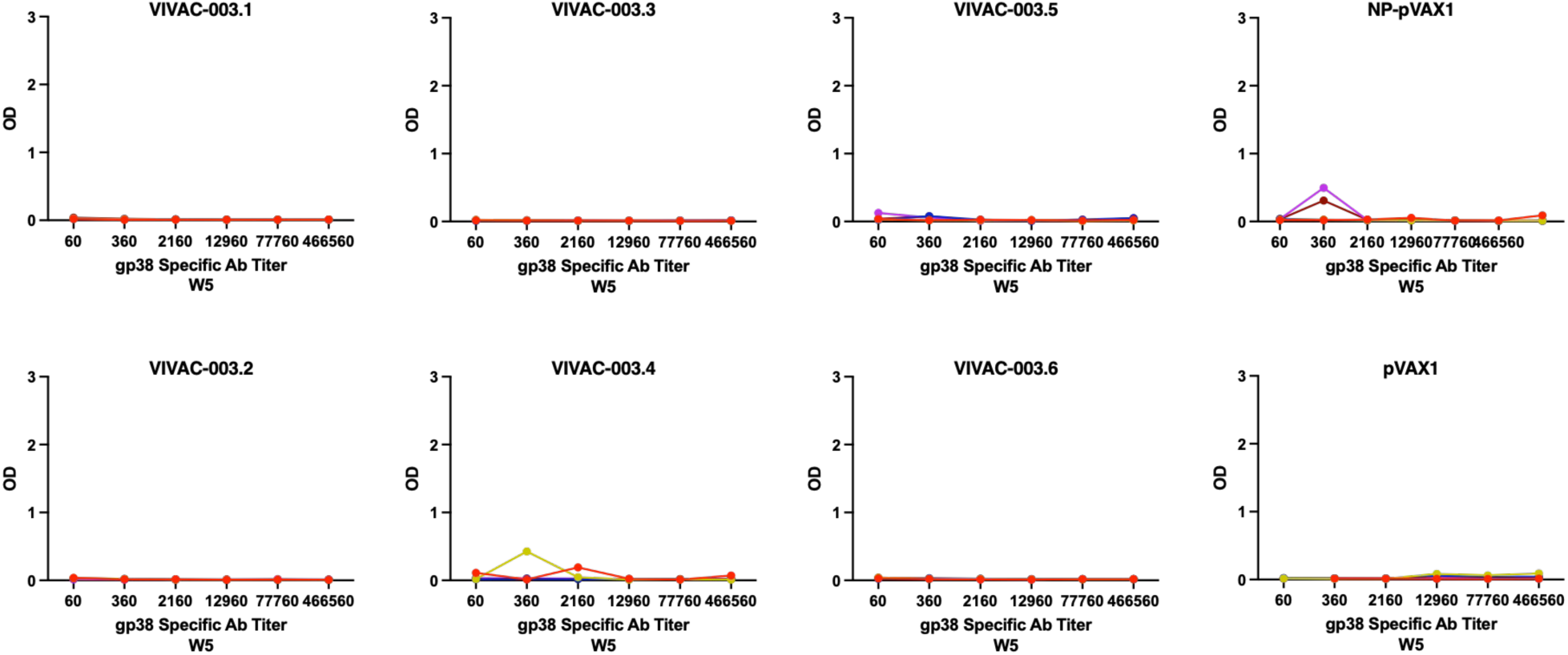
ELISA characterization of gp38-specific IgG responses across vaccine constructs five weeks post vaccination. gp38-specific serum IgG responses were quantified by ELISA using serially diluted sera collected from immunized mice. Optical density (OD) values were calculated as OD450–OD620 after background subtraction.

**Supplementary Figure S6.**
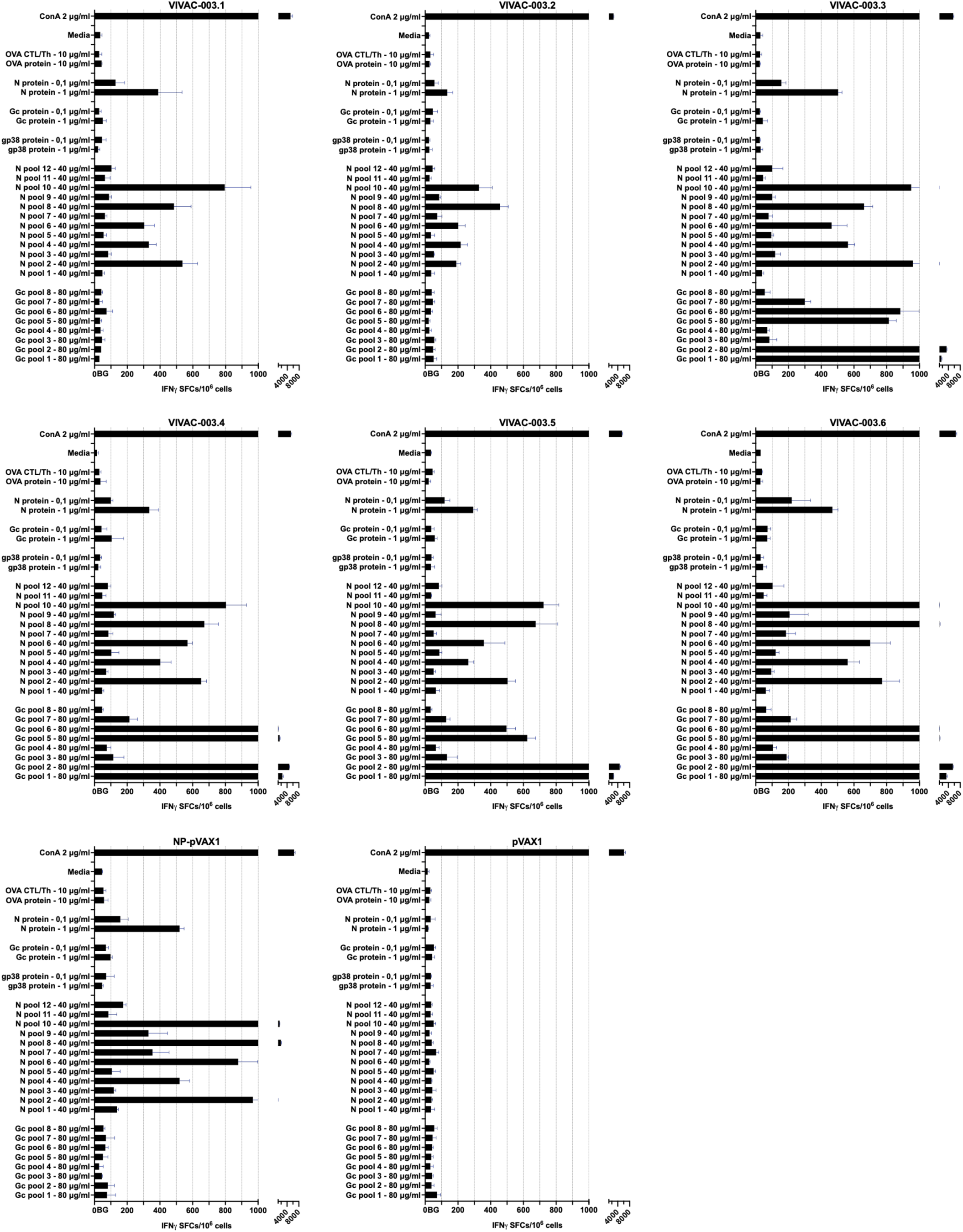
Individual group ELISpot results. IFN-γ–secreting cells were quantified by ELISpot and reported as spot-forming units (SFU) per 10⁶ splenocytes. Responses exceeding the mean of the medium-only background (BG) control were considered positive. Splenocytes were stimulated with Gc peptide pool (80 μg/ml), NP peptide pool (40 μg/ml), GP38 protein (1 μg/ml high, 0.1 μg/ml low), Gc protein (1 μg/ml high, 0.1 μg/ml low), N protein (1 μg/ml high, 0.1 μg/ml low), OVA protein (10 μg/ml), OVA CTL/Th peptides (10 μg/ml), or Concanavalin-A (ConA, 2 μg/ml). All stimulations were performed in triplicate, simple bars show mean and SD (n = 3).

**Supplementary Figure 7.**
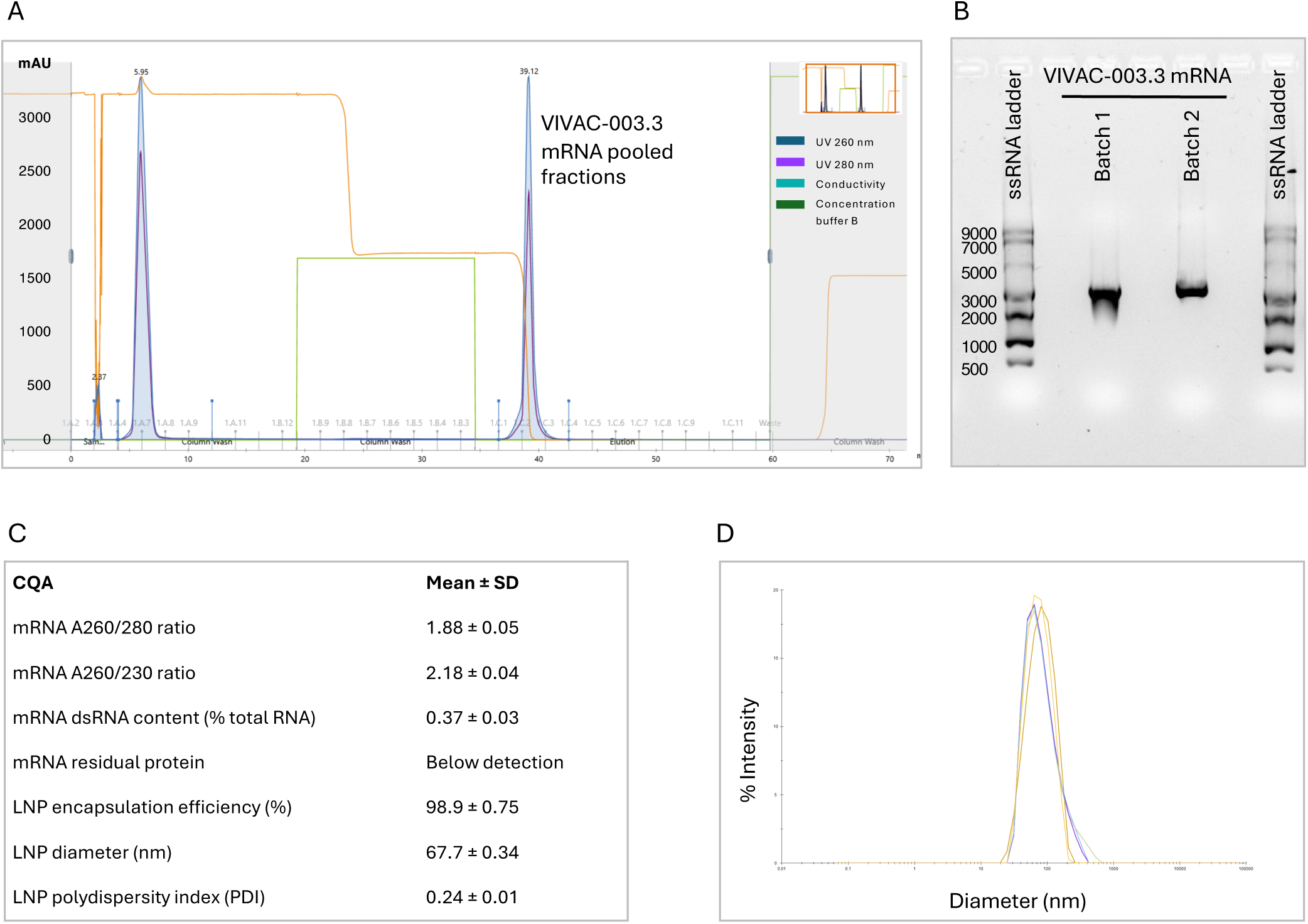
Quality control and characterization of independently manufactured SVF-003.3 mRNA-LNP batches. **(A)** Representative UV absorbance (260 nm) chromatogram from oligo(dT)-HPLC purification of IVT-produced SVF-003.3 mRNA. Full-length poly(A)-tailed mRNA was pooled from fractions C1–C3. **(B)** Agarose gel electrophoresis of purified full-length SVF-003.3 mRNA. An NEB single-stranded RNA (ssRNA) ladder was used as the molecular size reference. Samples were resolved on a 1% agarose gel at 160 V and visualized using GelRed staining. **(C)** Critical quality attributes (CQAs) of independently manufactured SVF-003.3 mRNA and mRNA-LNP batches. Values are presented as mean ± SD from two independently manufactured mRNA batches and four independently manufactured mRNA-LNP batches, respectively. dsRNA content (% of total RNA) was determined by the Lumit™ dsRNA Detection Assay, and residual protein by the AccuOrange assay, encapsulation efficiency (%) by the RiboGreen assay, particle size and polydispersity index (PDI) by dynamic light scattering (DLS). Residual protein was below the assay limit of detection. **(D)** Representative intensity-weighted particle size distribution measured by DLS. Data are shown from a single manufacturing batch and represent five replicate measurements, each comprising ten acquisitions.

**Supplementary Figure 8.**
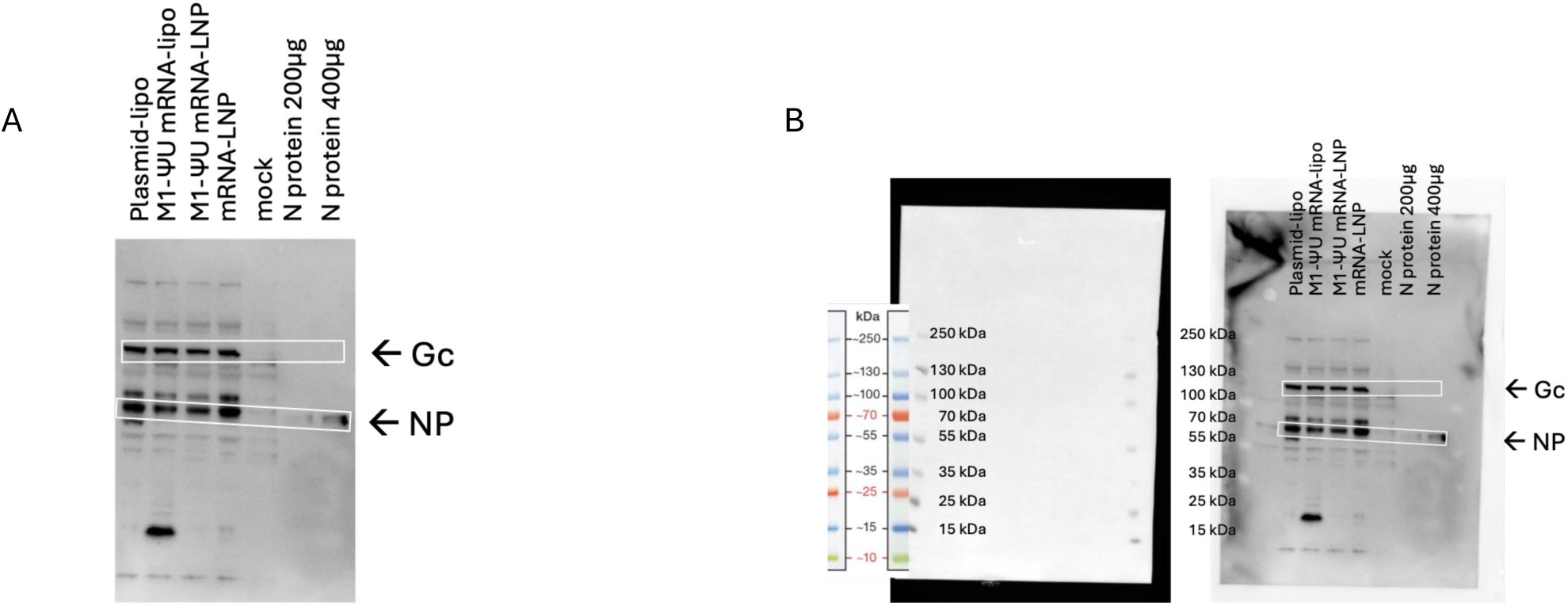
Western blot analysis of NP and Gc expression in transfected HEK2G3T cells. **A.** Cells were transfected with the indicated constructs, and lysates were collected 24 h after transfection. The NP or Gc protein was detected using immunized mice sera as primary antibody, untransfected cells were included as a negative control. Boxed regions indicate the expected target and loading-control bands. B. Uncropped images of the molecular-weight marker and the corresponding ECL-exposed blot are shown.

**Supplementary Figure S9.**
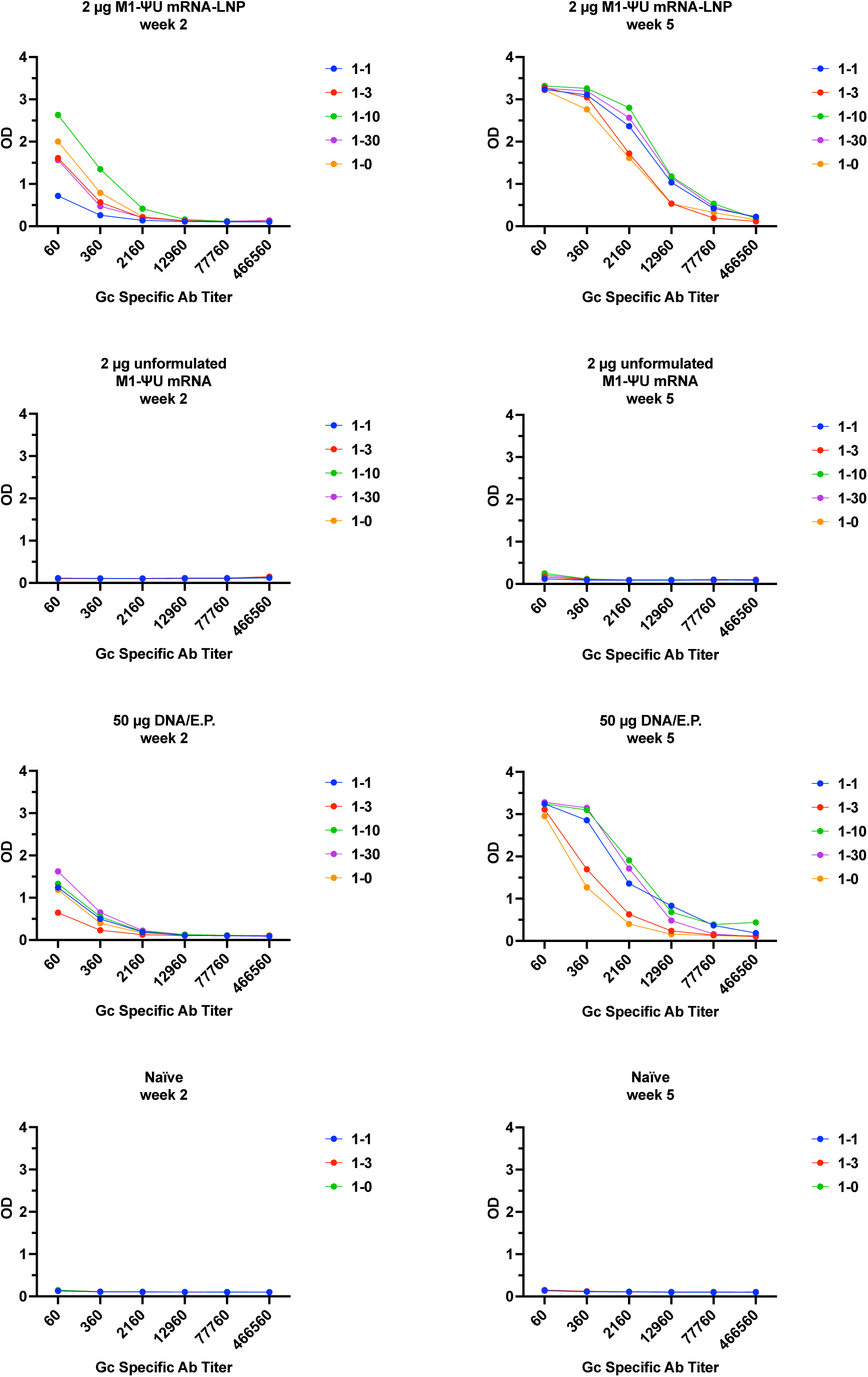
ELISA characterization of Gc-specific IgG responses across vaccine constructs post vaccination. Gc-specific serum IgG responses were quantified by ELISA using serially diluted sera collected from immunized mice. Optical density (OD) values were calculated as OD450–OD620 after background subtraction.

**Supplementary Figure S10.**
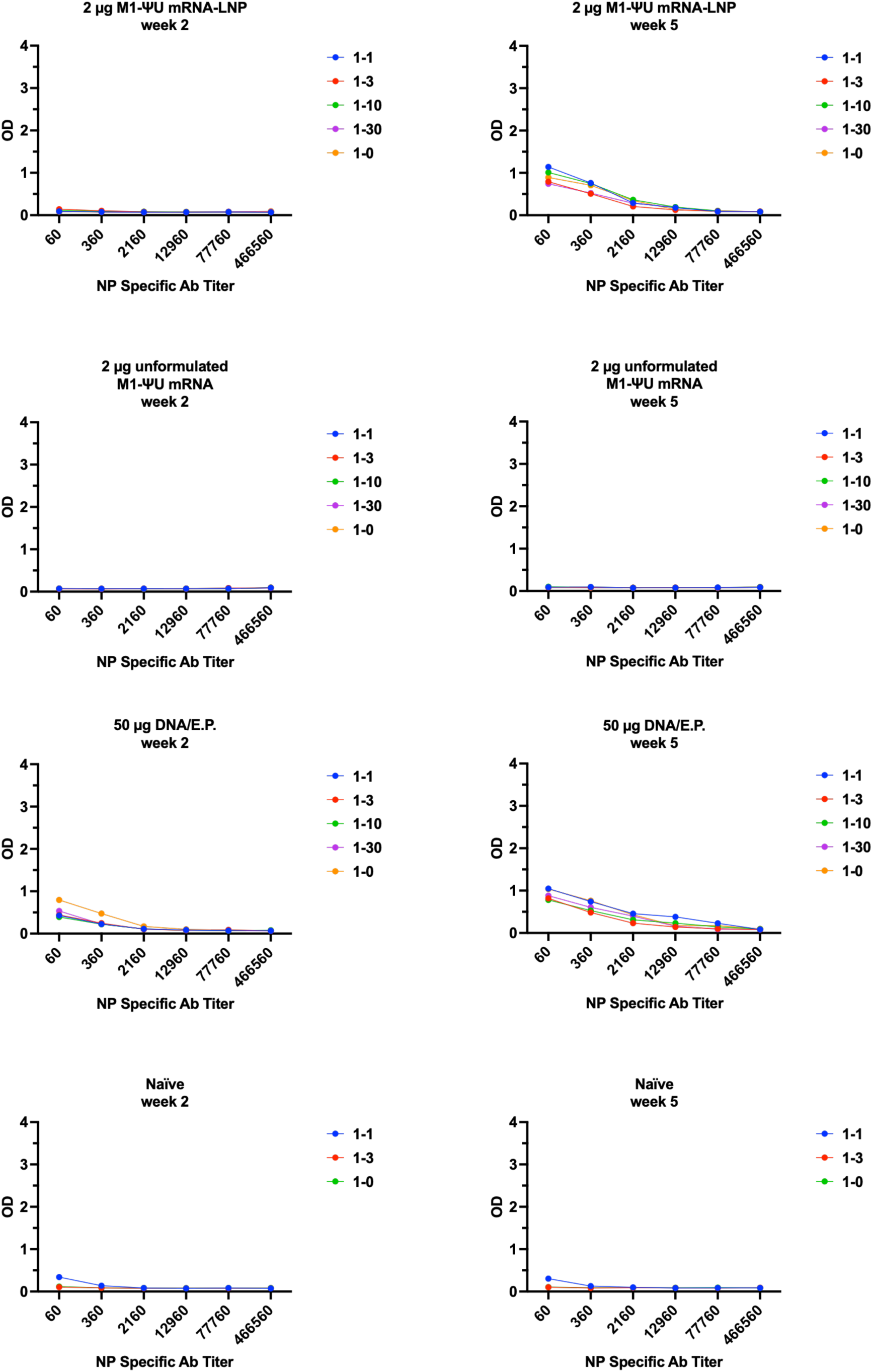
**ELISA characterization of NP-specific IgG responses across vaccine constructs post vaccination. NP**-specific serum IgG responses were quantified by ELISA using serially diluted sera collected from immunized mice. Optical density (OD) values were calculated as OD450–OD620 after background subtraction.

**Supplementary Figure S11.**
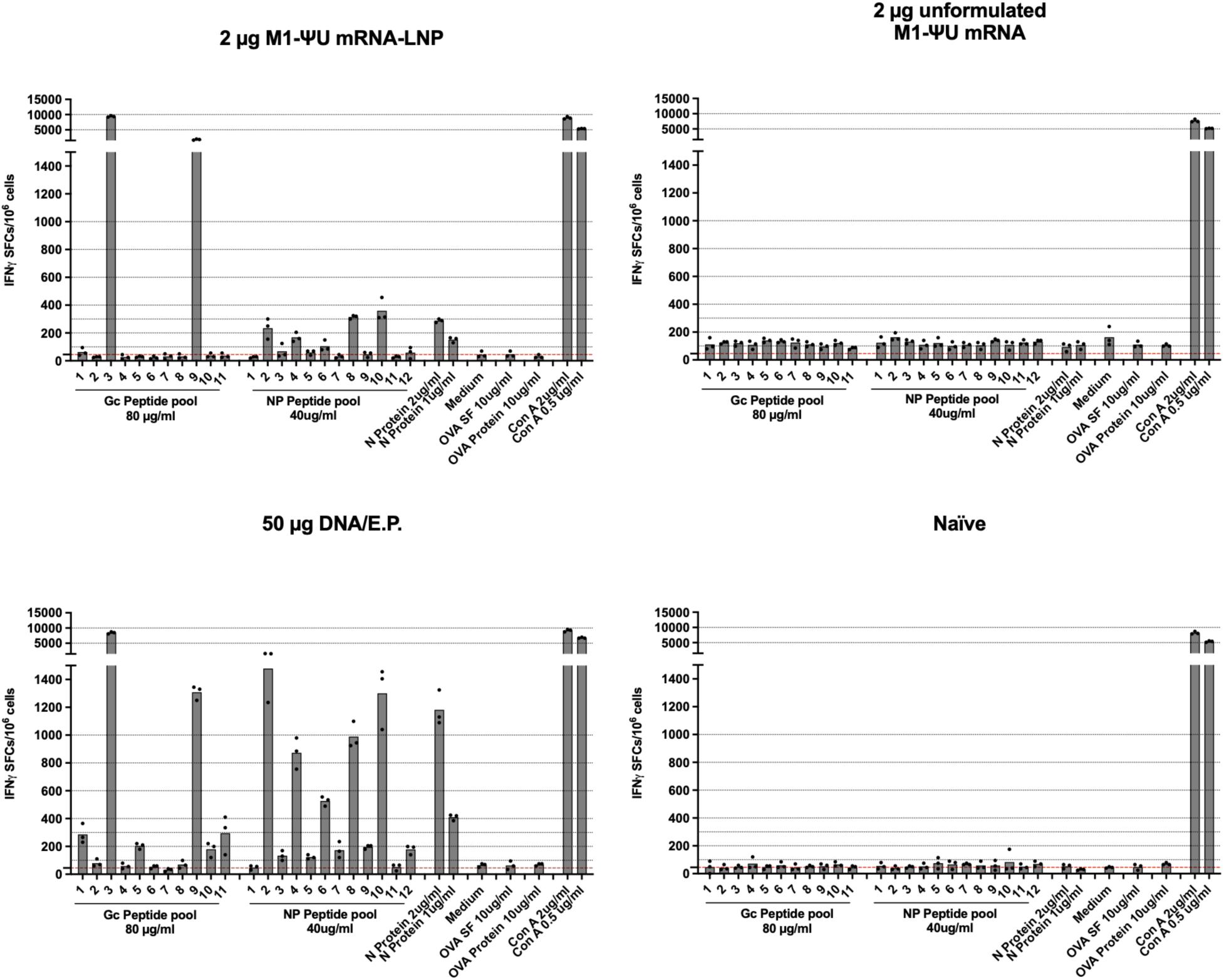
Individual group ELISpot results. IFN-γ–secreting cells were quantified by ELISpot and reported as spot-forming units (SFU) per 10⁶ splenocytes. Responses exceeding the mean of the medium-only background (BG) control were considered positive. Splenocytes were stimulated with Gc peptide pool (80 μg/ml), NP peptide pool (40 μg/ml), N protein (2 μg/ml and 1 μg/ml), OVA protein (10 μg/ml), OVA SF (10 μg/ml), or Concanavalin-A (ConA; 2 μg/ml and 0.5 μg/ml). All stimulations were performed in triplicate, symbol bars show mean and SD (n = 3).

**Supplementary Figure S12.**
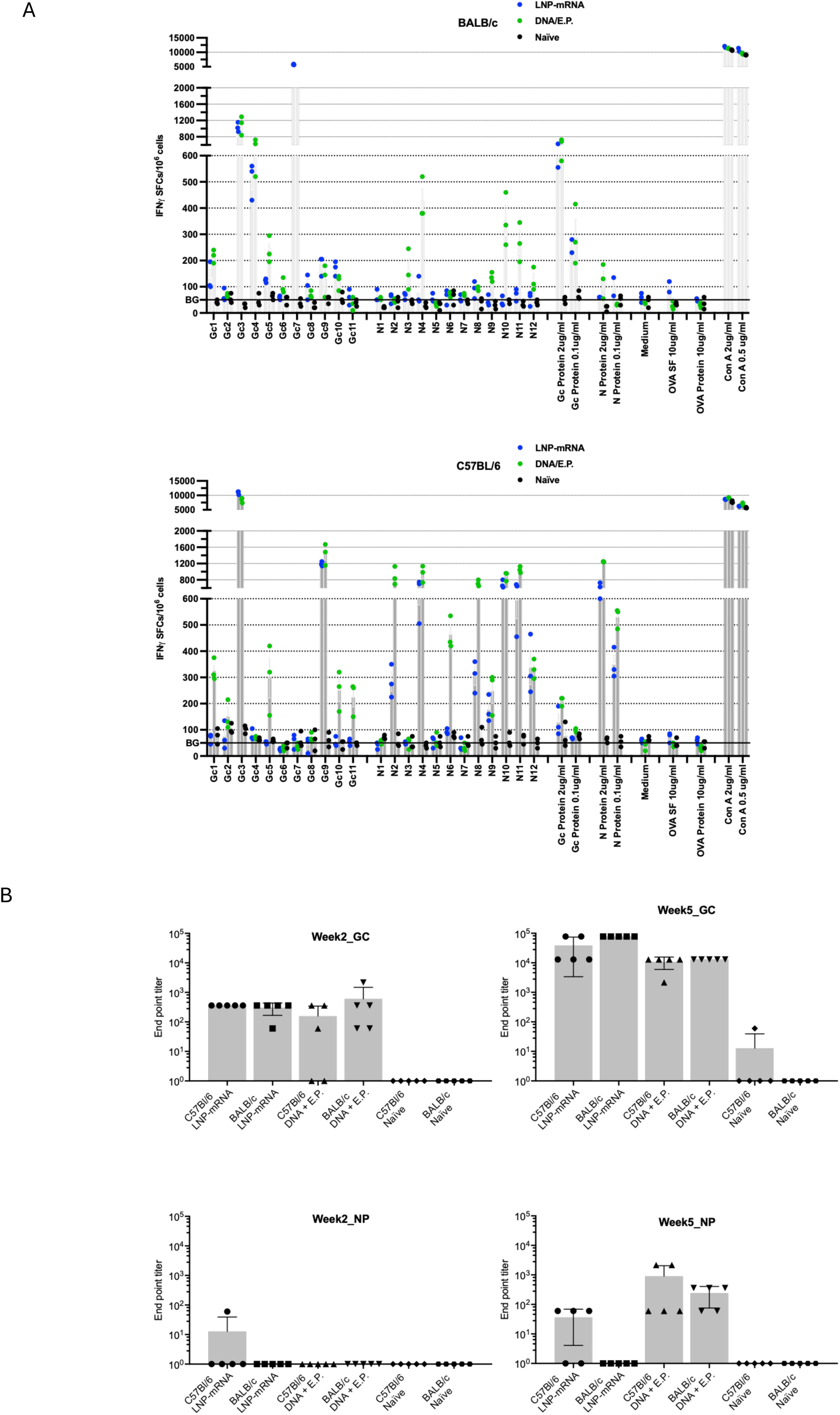
Comparison immunogenicty study performed in both BALB/c and C57BL/6J mice. **(A)** IFN-γ–secreting cells were quantified by ELISpot and reported as spot-forming units (SFU) per 10⁶ splenocytes. Responses exceeding the mean of the medium-only background (BG) control were considered positive. Splenocytes were stimulated with Gc peptide pool (80 μg/ml), NP peptide pool (40 μg/ml). All stimulations were performed in triplicate, simple bars show mean and SD (n = 3). **(B)** ELISA characterization of Gc-specific and N-specific IgG endpoint titers across vaccine constructs at week two and week five post vaccination.

**Supplementary Figure S13.**
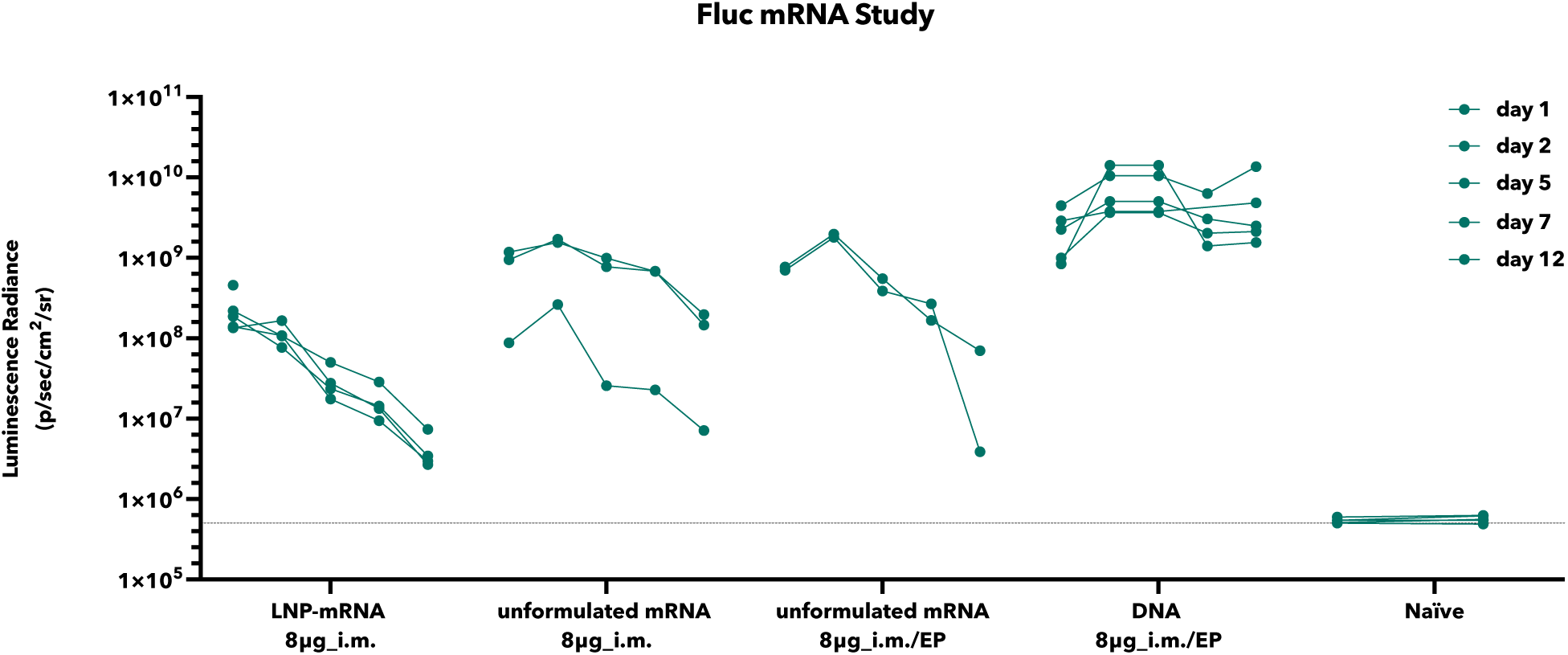
***In vivo* expression of luciferase following nucleic acid delivery assessed by IVIS imaging.** BALB/c (H-2^d^) mice (8–12 weeks old) were injected intramuscularly (i.m.) into the tibialis anterior (TA) muscle. Five experimental groups were included: 8 µg LNP-formulated mRNA, n=5, 8 µg unformulated mRNA, n=3, 8 µg unformulated mRNA delivered by electroporation (EP), n=2, 8 µg plasmid DNA delivered by EP, n=5, and a naïve control group receiving no injection, n=5. Bioluminescence imaging was performed using an *in vivo* imaging system (IVIS) on days 1, 2, 5, 7, and 12 post-administration to monitor luciferase expression at the injection site.

**Supplementary Figure S14.**
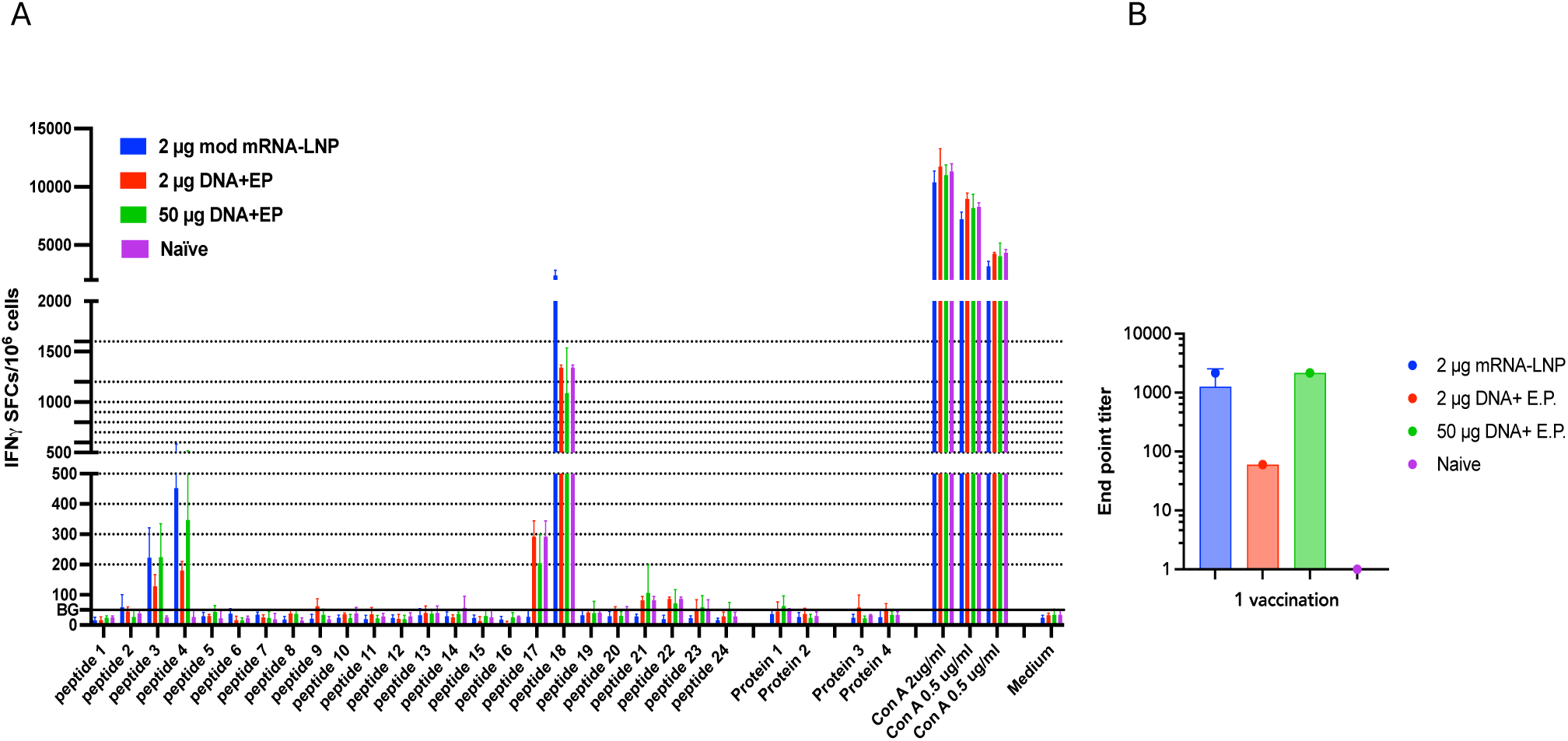
Immunogenicity comparison of mRNA and DNA vaccination. **A.** IFN-γ–secreting cells were quantified by ELISpot and reported as spot-forming units (SFU) per 10⁶ splenocytes. Responses exceeding the mean of the medium-only background (BG) control were considered positive. Splenocytes were stimulated with antigen-specific peptide pools or recombinant antigen proteins, with concanavalin A (ConA) included as a positive control. All stimulations were performed in triplicate, symbol bars show mean and SD (n = 3). **B.** Antigen-specific IgG responses were quantified by ELISA using serially diluted sera collected from immunized mice. Optical density (OD) values were calculated as OD450–OD620 following background subtraction.

## REFERENCE

1. Hawman, D. W. & Feldmann, H. Crimean-Congo haemorrhagic fever virus. Nat Rev Microbiol 21, 463–477 (2023).

2. Celina, S. S., Italiya, J., Tekkara, A. O. & Černý, J. Crimean-Congo haemorrhagic fever virus in ticks, domestic, and wild animals. Front. Vet. Sci. 11, (2025).

3. Atim, S. A. et al. Risk factors for Crimean-Congo Haemorrhagic Fever (CCHF) virus exposure in farming communities in Uganda. Journal of Infection 85, 693–701 (2022).

4. Çıtıl, R. et al. Determination of Seroprevalence and Risk Factors of Crimean–Congo Haemorrhagic Fever (CCHF) in the Endemic Region in Turkey: A Population-Based Cross-Sectional Study. J Trop Med 2021, 9945089 (2021).

5. Şahin, A. M., Ataş, E. & Çetin, S. Crimean-Congo hemorrhagic fever: Strategies for diagnosis at initial admission. The Brazilian Journal of Infectious Diseases 29, 104516 (2025).

6. Frank, M. G., Weaver, G. & Raabe, V. Crimean-Congo Hemorrhagic Fever Virus for Clinicians—Diagnosis, Clinical Management, and Therapeutics. Emerg Infect Dis 30, 864–873 (2024).

7. Frank, M. G., Weaver, G. & Raabe, V. Crimean-Congo Hemorrhagic Fever Virus for Clinicians—Virology, Pathogenesis, and Pathology. Emerg Infect Dis 30, 847–853 (2024).

8. Johnson, S. et al. Ribavirin for treating Crimean Congo haemorrhagic fever. Cochrane Database Syst Rev 6, CD012713 (2018).

9. Ozbey, S. B., Kader, Ç., Erbay, A. & Ergönül, Ö. Early use of ribavirin is beneficial in Crimean-Congo hemorrhagic fever. Vector Borne Zoonotic Dis 14, 300–302 (2014).

10. Muzammil, K. et al. Recent Advances in Crimean-Congo Hemorrhagic Fever Virus Detection, Treatment, and Vaccination: Overview of Current Status and Challenges. Biol Proced Online 26, 20 (2024).

11. Ahata, B. & Akçapınar, G. B. CCHFV vaccine development, current challenges, limitations, and future directions. Front Immunol 14, 1238882 (2023).

12. Ozdarendeli, A. Crimean–Congo Hemorrhagic Fever Virus: Progress in Vaccine Development. Diagnostics (Basel) 13, 2708 (2023).

13. Deyde, V. M., Khristova, M. L., Rollin, P. E., Ksiazek, T. G. & Nichol, S. T. Crimean-Congo Hemorrhagic Fever Virus Genomics and Global Diversity. Journal of Virology 80, 8834–8842 (2006).

14. Garrison, A. R. et al. Nucleocapsid protein-specific monoclonal antibodies protect mice against Crimean-Congo hemorrhagic fever virus. Nat Commun 15, 1722 (2024).

15. Leventhal, S. S. et al. Replicating RNA vaccination elicits an unexpected immune response that efficiently protects mice against lethal Crimean-Congo hemorrhagic fever virus challenge. eBioMedicine 82, 104188 (2022).

16. Rao, D. et al. CD8+ T-cells target the Crimean-Congo haemorrhagic fever virus Gc protein to control the infection in wild-type mice. eBioMedicine 97, 104839 (2023).

17. Fels, J. M. et al. Protective neutralizing antibodies from human survivors of Crimean-Congo hemorrhagic fever. Cell 184, 3486–3501.e21 (2021).

18. Mishra, A. K. et al. Structural basis of synergistic neutralization of Crimean-Congo hemorrhagic fever virus by human antibodies. Science 375, 104–109 (2022).

19. Appelberg, S. et al. Nucleoside-Modified mRNA Vaccines Protect IFNAR–/– Mice against Crimean-Congo Hemorrhagic Fever Virus Infection. J Virol 96, e01568–21 (2022).

20. Duh, D. et al. The complete genome sequence of a Crimean-Congo Hemorrhagic Fever virus isolated from an endemic region in Kosovo. Virol J 5, 7 (2008).

21. Haddock, E. et al. A cynomolgus macaque model for Crimean-Congo haemorrhagic fever. Nat Microbiol 3, 556–562 (2018).

22. Hawman, D. W. et al. Immunocompetent mouse model for Crimean-Congo hemorrhagic fever virus. Elife 10, e63906 (2021).

23. Leventhal, S. S. et al. Replicating RNA vaccine confers durable immunity against Crimean Congo hemorrhagic fever virus challenge in mice. NPJ Vaccines 9, 249 (2024).

24. Karaaslan, E. et al. Crimean Congo hemorrhagic fever virus nucleoprotein and GP38 subunit vaccine combination prevents morbidity in mice. npj Vaccines 9, 148 (2024).

25. Suschak, J. J. et al. A CCHFV DNA vaccine protects against heterologous challenge and establishes GP38 as immunorelevant in mice. npj Vaccines 6, 31 (2021).

26. Hawman, D. W. et al. A DNA-based vaccine protects against Crimean-Congo haemorrhagic fever virus disease in a Cynomolgus macaque model. Nat Microbiol 6, 187–195 (2021).

27. Tipih, T. et al. Single dose VSV-based vaccine protects mice against lethal heterologous Crimean-Congo hemorrhagic fever virus challenge. npj Vaccines 10, 109 (2025).

28. Leventhal, S. S. et al. Antibodies targeting the Crimean-Congo Hemorrhagic Fever Virus nucleoprotein protect via TRIM21. Nat Commun 15, 9236 (2024).

29. Rao, J. et al. A single-dose mRNA vaccine protects mice from lethal Crimean-Congo hemorrhagic fever virus infection. npj Vaccines 10.1038/s41541-025-01310-x (2025) doi:10.1038/s41541-025-01310-x.

30. Leventhal, S. S. et al. Single dose, dual antigen RNA vaccines protect against lethal Crimean-Congo haemorrhagic fever virus infection in mice. eBioMedicine 101, 105017 (2024).

31. Golden, J. W. et al. GP38-targeting monoclonal antibodies protect adult mice against lethal Crimean-Congo hemorrhagic fever virus infection. Science Advances 5, eaaw9535 (2019).

32. Liu, M. A. A Comparison of Plasmid DNA and mRNA as Vaccine Technologies. Vaccines (Basel) 7, 37 (2019).

33. Sardesai, N. Y. & Weiner, D. B. Electroporation Delivery of DNA Vaccines: Prospects for Success. Curr Opin Immunol 23, 421–429 (2011).

34. Vasan, S. et al. In Vivo Electroporation Enhances the Immunogenicity of an HIV-1 DNA Vaccine Candidate in Healthy Volunteers. PLOS ONE 6, e19252 (2011).

35. Kisakov, D. N. et al. The use of electroporation to deliver DNA-based vaccines. Expert Review of Vaccines 23, 102–123 (2024).

36. Xia, D. et al. An ultra-low-cost electroporator with microneedle electrodes (ePatch) for SARS-CoV-2 vaccination. Proceedings of the National Academy of Sciences 118, e2110817118 (2021).

37. Pardi, N., Hogan, M. J., Porter, F. W. & Weissman, D. mRNA vaccines — a new era in vaccinology. Nat Rev Drug Discov 17, 261–279 (2018).

38. Alameh, M.-G. et al. Lipid nanoparticles enhance the efficacy of mRNA and protein subunit vaccines by inducing robust T follicular helper cell and humoral responses. Immunity 54, 2877–2892.e7 (2021).

39. Andrade, V. M. et al. Delineation of DNA and mRNA COVID-19 vaccine-induced immune responses in preclinical animal models. Hum Vaccin Immunother 19, 2281733.

40. Aleman, S. et al. A phase 1 clinical trial of a multi-antigen SARS-CoV DNA vaccine as a booster after dose three spike-based mRNA vaccinations. Molecular Therapy Advances 34, 201710 (2026).

41. Blakney, A. K. et al. Safety and efficacy of mRNA vaccines: a mechanistic and public health perspective. The Lancet 10.1016/S0140-6736(26)00512-X (2026) doi:10.1016/S0140-6736(26)00512-X.

42. Connors, J. et al. Lipid nanoparticles (LNP) induce activation and maturation of antigen presenting cells in young and aged individuals. Commun Biol 6, 188 (2023).

43. Ndeupen, S. et al. The mRNA-LNP platform’s lipid nanoparticle component used in preclinical vaccine studies is highly inflammatory. *bioRxiv* 2021.03.04.430128 (2021) doi:10.1101/2021.03.04.430128.

44. Kozma, G. T. et al. Role of anti-polyethylene glycol (PEG) antibodies in the allergic reactions to PEG-containing Covid-19 vaccines: Evidence for immunogenicity of PEG. Vaccine 41, 4561–4570 (2023).

45. Zeng, C., Zhang, C., Walker, P. G. & Dong, Y. Formulation and Delivery Technologies for mRNA Vaccines. Curr Top Microbiol Immunol 440, 71–110 (2022).

46. 46. Guidance for Industry: Considerations for Plasmid DNA Vaccines for Infectious Disease Indications. Biotechnology Law Report 26, 641–648 (2007).

47. Patel, A. et al. Intradermal-delivered DNA vaccine induces durable immunity mediating a reduction in viral load in a rhesus macaque SARS-CoV-2 challenge model. Cell Reports Medicine 2, 100420 (2021).

48. Kulkarni, V. et al. DNA vaccination by intradermal electroporation induces long-lasting immune responses in rhesus macaques. J Med Primatol 43, 329–340 (2014).

49. Brunner, W. M. et al. Comparison of antibody response durability of mRNA-1273, BNT162b2, and Ad26.COV2.S SARS-CoV-2 vaccines in healthcare workers. Int J Infect Dis 123, 183–191 (2022).

50. Yamamoto, S. et al. Durability and determinants of anti-SARS-CoV-2 spike antibodies following the second and third doses of mRNA COVID-19 vaccine. Clinical Microbiology and Infection 29, 1201.e1–1201.e5 (2023).

51. Trimble, C. L. et al. Safety, efficacy, and immunogenicity of VGX-3100, a therapeutic synthetic DNA vaccine targeting human papillomavirus 16 and 18 E6 and E7 proteins for cervical intraepithelial neoplasia 2/3: a randomised, double-blind, placebo-controlled phase 2b trial. The Lancet 386, 2078–2088 (2015).

52. Schmeer, M., Buchholz, T. & Schleef, M. Plasmid DNA Manufacturing for Indirect and Direct Clinical Applications. Human Gene Therapy 28, 856–861 (2017).

53. Buckland, B. et al. Vaccine process technology—A decade of progress. Biotechnology and Bioengineering 121, 2604–2635 (2024).

54. Whitley, J. et al. Development of mRNA manufacturing for vaccines and therapeutics: mRNA platform requirements and development of a scalable production process to support early phase clinical trials. Transl Res 242, 38–55 (2022).

55. Yan, J. et al. Distinct roles of vaccine-induced SARS-CoV-2-specific neutralizing antibodies and T cells in protection and disease. Mol Ther 32, 540–555 (2024).

56. Appelberg, S. et al. A universal SARS-CoV DNA vaccine inducing highly cross-reactive neutralizing antibodies and T cells. EMBO Mol Med 14, e15821 (2022).

57. Devignot, S., Bergeron, E., Nichol, S., Mirazimi, A. & Weber, F. A Virus-Like Particle System Identifies the Endonuclease Domain of Crimean-Congo Hemorrhagic Fever Virus. J Virol 89, 5957–5967 (2015).

58. Müller, M. A. et al. Evidence for widespread infection of African bats with Crimean-Congo hemorrhagic fever-like viruses. Sci Rep 6, 26637 (2016).

